# 3D osteocyte networks under Pulsatile Unidirectional Fluid Flow Stimuli (PUFFS)

**DOI:** 10.1101/2025.04.30.651549

**Authors:** Anna-Blessing Merife, Arun Poudel, Angelika Polshikova, Zachary Geffert, Jason A. Horton, Mohammad Mehedi Hasan Akash, Anupum Pandey, Saikat Basu, Daniel Fougnier, Pranav Soman

**Affiliations:** Department of Chemical and Biomedical Engineering, L.C. Smith College of Engineering Syracuse University, Syracuse, NY 13244 USA; Department of Neuroscience and Physiology, Alan and Marlene Norton College of Medicine, SUNY Upstate Medical University, Syracuse NY 13210 USA; Department of Mechanical Engineering, South Dakota State University, Brookings, SD 57007, United States; Department of Mechanical and Aerospace Engineering, L.C. Smith College of Engineering Syracuse University, Syracuse, NY 13244 USA

**Keywords:** Osteocytes, 3D cultures, MLO-Y4, mechanical stimuli, in vitro model, microfluidic, chip

## Abstract

Although osteocytes are known to play a key role in skeletal mechano-adaptation, few in vitro models have investigated how pulsatile mechanical stimuli influence the properties of 3D osteocyte networks. Here we design and develop a microfluidic based *in vitro* model to study 3D osteocyte networks cultured under Pulsatile Unidirectional Fluid Flow Stimuli (PUFFS). Digital light projection stereolithography was used to design and fabricate a three-chambered PDMS microfluidic chip. Model osteocytes (murine MLO-Y4) were encapsulated in collagen matrix within the chip to form self-assembled three-dimensional (3D) cell networks. Daily stimulus in the form of PUFFS was then applied for upto 21 days. A combination of experiments, computational simulation and analytical modeling was used to characterize the mechanical environment experienced by embedded cells during PUFFS. Viability, morphology, cell-connectivity, expression of key proteins, and gene expression, and real-time calcium signaling within 3D osteocyte networks were characterized at select time-points and compared to static conditions. Results show that PUFFS stimulation at 0.33 and 1.66 Hz can initiate mechanotransduction via calcium signals that are propagated across the network of collagen encapsulated osteocytes via Cx43 junctions. Furthermore, osteocytes cultured in these devices maintain expression of several key osteocyte genes for up to 21d. Taken together, this model can potentially serve as a testbed to study how 3D osteocyte networks respond to dynamic mechanical stimulation relevant to skeletal tissues.

## INTRODUCTION

Osteocytes are the primary mechanosensory cells within bone tissue. Mechanical loading creates interstitial fluid flow that induces dynamic signaling across three-dimensional (3D) networks of interconnected osteocytes. In turn, these signals spatially coordinate osteoblastic bone deposition and osteoclastic resorption at the bone surface via paracrine and juxtacrine factors.^1–9^ Mechanical stimulation is necessary for osteocyte function^10^, and disruption of their mechanotransduction is implicated in many skeletal disorders.^11–16^ Contemporary in vivo and ex vivo models have yet to reveal the mechanisms that propagate short-term signals such as calcium across 3D networks, that are modulate long-term remodeling responses.^11–22^ This is largely due to the dependence on live animals or human explant tissue^23–26^, which require expensive and complicated experimental apparatus^27–29^ with low throughput and poor reproducibility and superficial depth of observation. To address these challenges, many complimentary in vitro models have been developed, although most studies continue to rely on simpler two-dimensional (2D) designs that do not replicate the complex 3D architecture and dynamic signaling of osteocyte networks *in vivo*. For example, bulk stimulation of osteocyte monolayers via flow chambers subjects nearly all cells in the system to identical and simultaneous stimuli^30–32^. More sophisticated tools such as nanoindentation can stimulate individual osteocytes within patterned 2D networks^33–37^ yet this does not consider the 3D microenvironment. Transwell models and microfluidic devices^38–48^ have been used to study paracrine signaling in a co-culture setup, however applying regionally confined mechanical stimulation especially during long-term cultures remains challenging. Pseudo-3D network models have also been developed by culturing osteocytes on the surfaces of 3D microbeads,^49^ or embedding cells within mineralized 3D constructs to mimic *in vivo* microenvironmental conditions,^48^ however their opacity precludes real-time visualization of signaling behavior. As a result, cells encapsulated within collagen continue to be the gold-standard to study real-time signaling within 3D osteocyte networks.^49–60^ However, new models need to be developed that allow (i) 3D osteocyte culture, (ii) application of defined mechanical stimuli, (iii) longitudinal study of real-time signal propagation between interconnected cells, and (iv) long term changes in osteocyte morphology, viability, proliferation, and gene expression. In this work, we report on the design, development and characterization of such a model to study how 3D osteocyte networks respond to dynamic stimuli. We developed and optimized apparatus to apply Pulsed Unidirectional Fluid Flow Stimuli (PUFFS) to 3D osteocyte-laden collagen networks and reliably capture calcium signaling by real-time fluorescence microscopy. Using a combination of empirical experiments, modeling and simulation approaches, we characterized the force microenvironment experienced by osteocytes in 3D networks during PUFFS. Lastly, we demonstrate that PUFFS can be applied to the 3D osteocyte networks for at least 21 days, allowing long-term assessment of changes in cell viability, morphology, connectivity, gene expression, and real-time signaling dynamics in response to stimulation.

## RESULTS AND DISCUSSION

### Design and fabrication of multi-chambered microfluidic chips

For long-term culture of 3D osteocyte networks, we designed and developed three-chambered microfluidic chips in PDMS. **(Fig.1A)** Briefly, Digital Light Projection (DLP) stereolithography was used to print a negative master mold using polyethylene glycol diacrylate (PEGDA) resin followed by replica-casting using PDMS and irreversibly bonding the PDMS molds to glass coverslips (22mm x 22mm, **Fig.1B**); the details of this process are explained in the Methods section. The final devices consist of three-chambers with inlet and outlet ports (2 mm diameter), a central chamber (850 μm wide, to house osteocyte-laden collagen) flanked on either side by two chambers (~500 μm wide, for PUFFS stimuli and media exchange), separated by an array of posts with an inter-post gap of 65 μm. **(Fig.1Bi)** The height of all the chambers within the chip was 250 µm. Post-fabrication, chips were surface coated with polydopamine (PDA) and MLO-Y4s in type I collagen (2.5 mg/mL)^61^ at a final concentration of 1×10^5^ cells/mL was thermally crosslinked (370C, 30 min) within the central chamber of the chip (Ch#2). The post array prevents leakage of cell-solution into side chambers during the gelation of collagen in chamber 2. For dynamic conditions, PUFFS (0.33, 15mins/daily; from Day 3-21) was applied to chamber 3 (Ch#3) of the chips while for static conditions no stimuli were applied. **(Fig.1Bii)**

### Setup design, development and optimization for PUFFS

To generate cyclic mechanical stimuli, we designed a new experimental setup that includes a peristaltic pump, controller, and connector tubings to generate pulsed unidirectional fluid flow stimuli or PUFFS applied at a frequency setting of 0.33 Hz (or 1.66Hz) in chamber #3 of the microfluidic chips. Pump pressure of 30 kPa results in a velocity of 0.018 m/s in chamber #3 of the chip; these experimental conditions do not cause any disruptions to the crosslinked collagen barrier for the duration of the study. **Figure 2A** shows the setup used to apply PUFFS to 3 independent chips. Next, we tested the ability of this setup to reliably capture calcium responses of 3D osteocyte culture during PUFFS. After gelation of MLO-Y4 laden collagen in chamber 2 of the chips, Fluo-4AM calcium dye was incubated and washed, and PUFFS was applied in Ch#3 for 60 seconds. The first setup coined as ‘open-loop’ involved an unidirectional flow with outlet tubing being open. Representative plots of calcium signaling in individual MLO-Y4s during PUFFS showed unstable profile; this is potentially due to the negative pressure and associated flow fluctuations due to backflows. **(Fig.2Ci)** To improve this setup, we tested a ‘closed-loop’ setup where both inlet and outlet tubings were used to generate a recycled unidirectional flow. This resulted in more stable signals but slight movement of the chip during the application of PUFFS caused fluctuations in the signal. To further improve reprocibility of signals, a stabilizer was designed and 3D printed to mitigate unwarranted movement from the connected chips during imaging; details of the stabilizer are provided in **Fig.2B, SI-1**. Since use of stabilizer and running PUFFS under closed-loop conditions provided reproducible calcium signal recordings, this setup was used for all experiments in this work.

**Figure 1.**
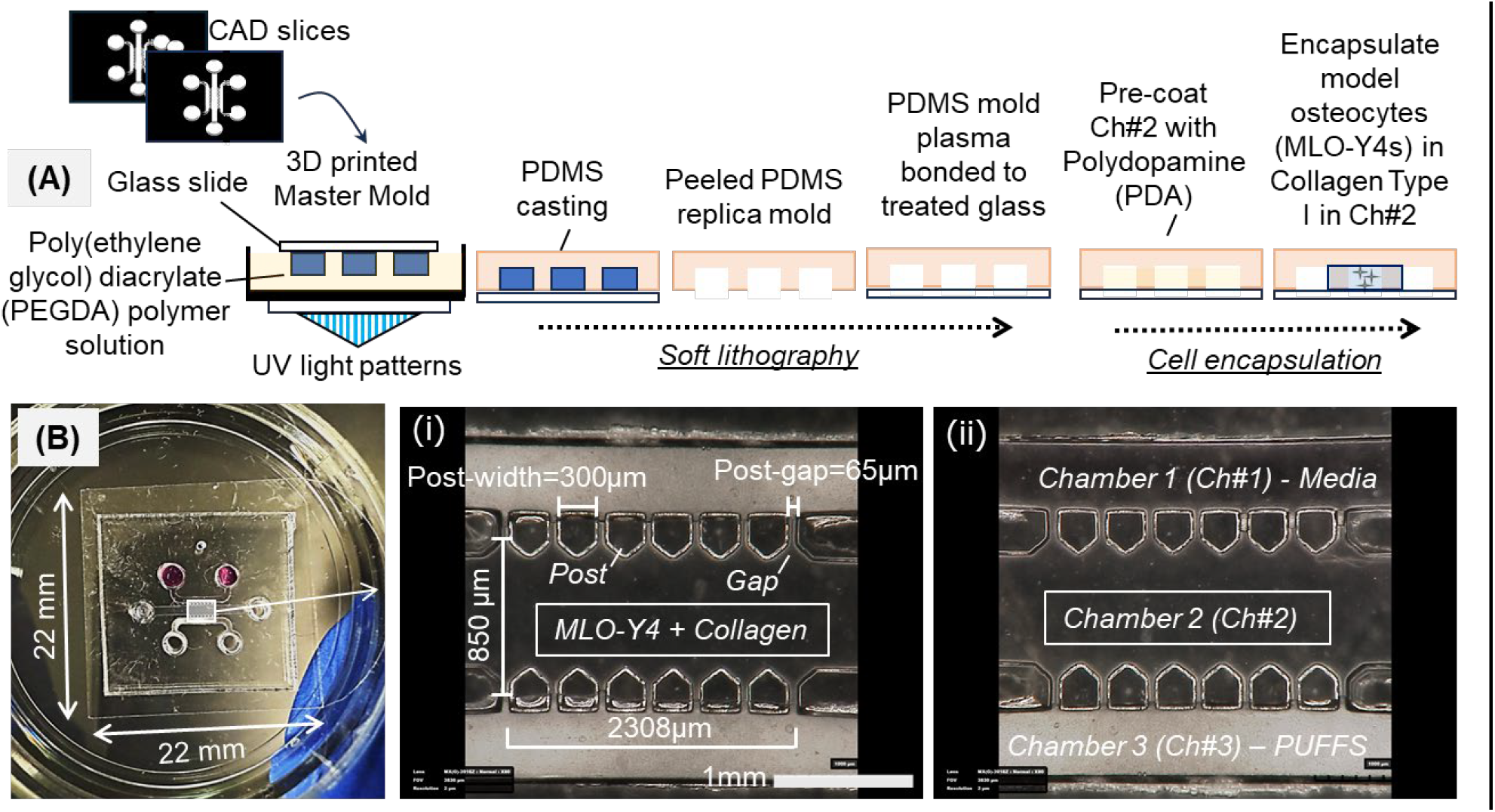
(A) Fabrication process flow to develop PDMS microfluidic chips with MLO-Y4-laden collagen barrier in Ch#2. (B) Representative picture of three-chambered chip showing three inlet-outlet pairs. (i, ii) Pictures shows relevant dimensions of chambers 1, 2 and 3 (Ch#1, #2, #3). Ch#2 house MLO-Y4 + collagen gel. For dynamic culture, PUFFS is applied in Ch#3 with static media culture in Ch#1. For static culture, media is present in both Ch#1 and Ch#3.

**Figure 2.**
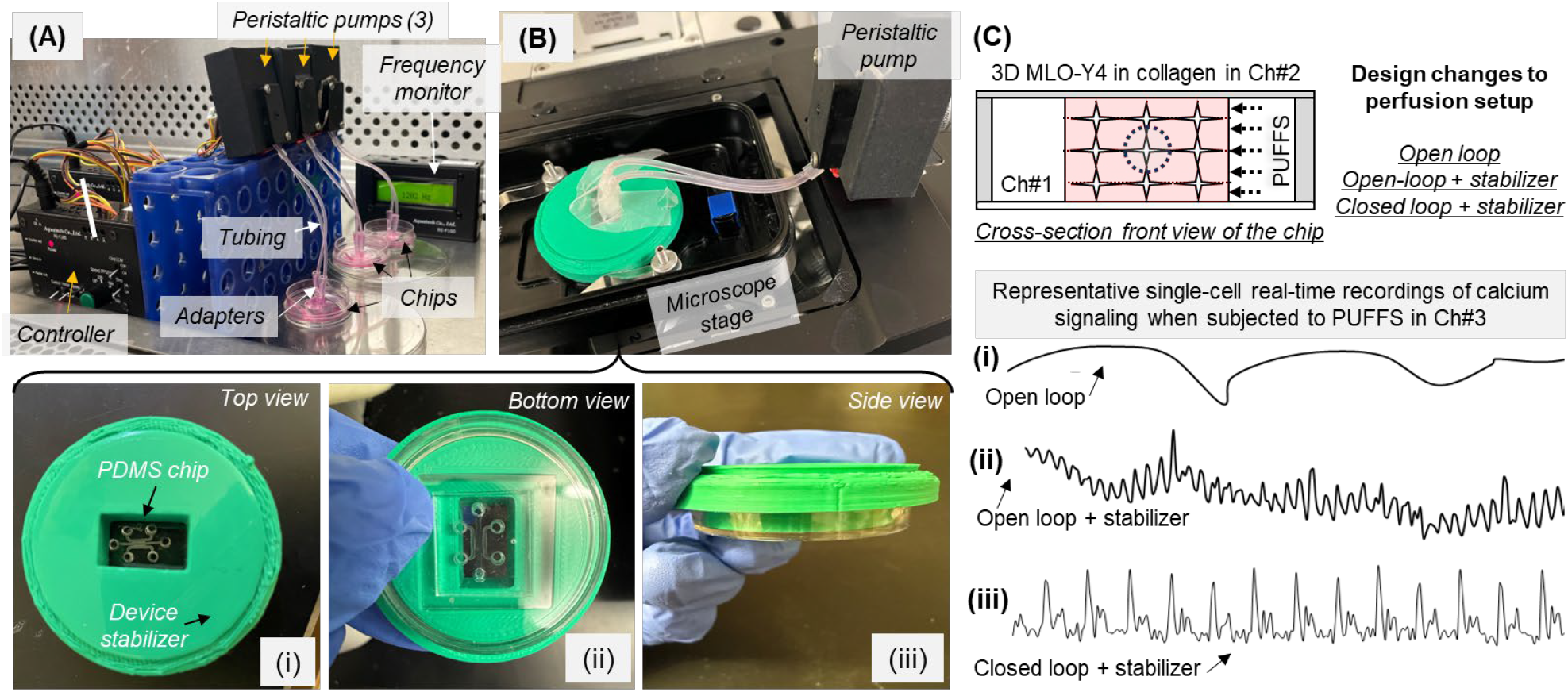
(A) Picture of setup showing parallel application of PUFFS to 3 chips. (B) Device stabilizer (shown in green) fitted onto microscope stage holds the chips and enables reproducible collection of real-time calcium signaling data during the application of PUFFS in Ch#3; (i-iii) show various views of the assembly of the chip within the stabilizer. (C) Schematic showing the cross-sectional view of the chip; (i-iii) shows changes in single-cell calcium signaling during PUFFS under different setup configurations.

### Characterization of mechanical microenvironment experienced by cells during PUFFS

Before biological characterization of osteocyte-laden collagen, it was important to understand the stresses experienced within collagen gel during PUFFS. For the collagen concentration used in this work (2.5 mg/ml), the storage and loss modulus were calculated as 159±33.51 Pa and 53.33±9.6 Pa, respectively **(SI-2)**. To characterize the stress and velocity profiles in crosslinked collagen when subjected to PUFFS, we used a combination of experiments, simulation, and modeling. First, collagen solution (2.5 mg/ml) was mixed with fluorescent beads (1 µm diameter) and crosslinked within chamber #2 of the chip. Upon application of PUFFS (0.33 Hz), collagen gel compresses and relaxes with each fluid flow pulse. **(Fig.3A i, ii, Video-1)** Although beads encapsulated within collagen aggregate into clusters (~25 µm), their displacements in various regions within collagen was used to generate a vector field map; here length of blue arrow indicated the magnitude of displacement **(Fig.3Aiii)**. Results show bead displacement of 46.8 µm in collagen sub-region that is proximal to PUFFS and 32.5 µm in those that are distal to PUFFS. Based on chip dimensions, we calculated a maximum shear stress of 56.17 kPa at the collagen surface (interface of Ch#2 and Ch#3). To calculate the stress within collagen located in chamber 2 (Ch#2), we developed an elasticity model **(Fig.2Bi, ii)** where we approximate the shear stress as normal loading (*p*_*c*_ in **Figure 3Ci** acting perpendicular to the collagen surface in Ch#3). This approximation is motivated by the observation that measured displacement in the collagen layer is predominantly in the direction normal to the interface. In this model, we assume the collagen layer to be purely elastic, and normal loading between two adjacent posts with the layer thickness (2h) representing the width of collagen in chamber 2. **Figure 3Biii** shows the distribution of compressive stress in collagen (*σ*_*yy*_) during PUFFS. Note that, during PUFFS, the stress is maximum in chamber 3 (collagen region between two adjacent microposts) reaching a value of 1 (blue color in **Fig.3Cii**) while regions behind the posts remain relatively stress free (marked by the red color in **Fig.3Cii**). This is the reason we choose to only analyze calcium signaling of MLO-Y4s located between the posts in this study – the region that experiences stresses during PUFFS. Three cross-sections marked by red, dashed lines in **Figure 3Bii**, were used to assess how the compressive stress (σyy) varies in collagen sub-regions proximal, central, and distal to PUFFS. Taking the highest value of 57.16kPa (*p*_*c*_) experienced by the collagen at the interface of Ch#3 and Ch#2, sub-regions proximal to PUFFS (0-285 μm) will experience stress of 57.16 – 45.16 kPa, central sub-region (286-570 µm) will experience stress of 45.16 – 22.47 kPa, while distal sub-regions (571-855 μm) will experience a stress range of 22.47 – 11.23 kPa. **(Fig.3Biv)** Lastly, to simulate the velocity distribution within the collagen gel during PUFFS, we developed a Eulerian viscous two-phase model, treating water as the primary phase (Ch#3) and the collagen gel as the secondary phase (Ch#2). **(Fig.3Bv)** The velocity contour plot in the collagen bulk reveals a high-velocity region at the collagen interface between Ch#3 and Ch#2 and at the edges of the posts and decline in velocity within the bulk (Ch#2) due to collagen’s viscous resistance. By comparing the experimentally measured bead velocities at specific points within the collagen to the simulated values, we identified correction factors that aligned the numerical predictions with the experimental data, thereby validating our simulation model. Results show that the spatial demarcations of the simulated flow barriers in the collagen (Ch#2) match well with the experimental trends. Details related to experimental calculations, analytical modeling, and simulations can be found in **SI.3-5**.

**Figure 3.**
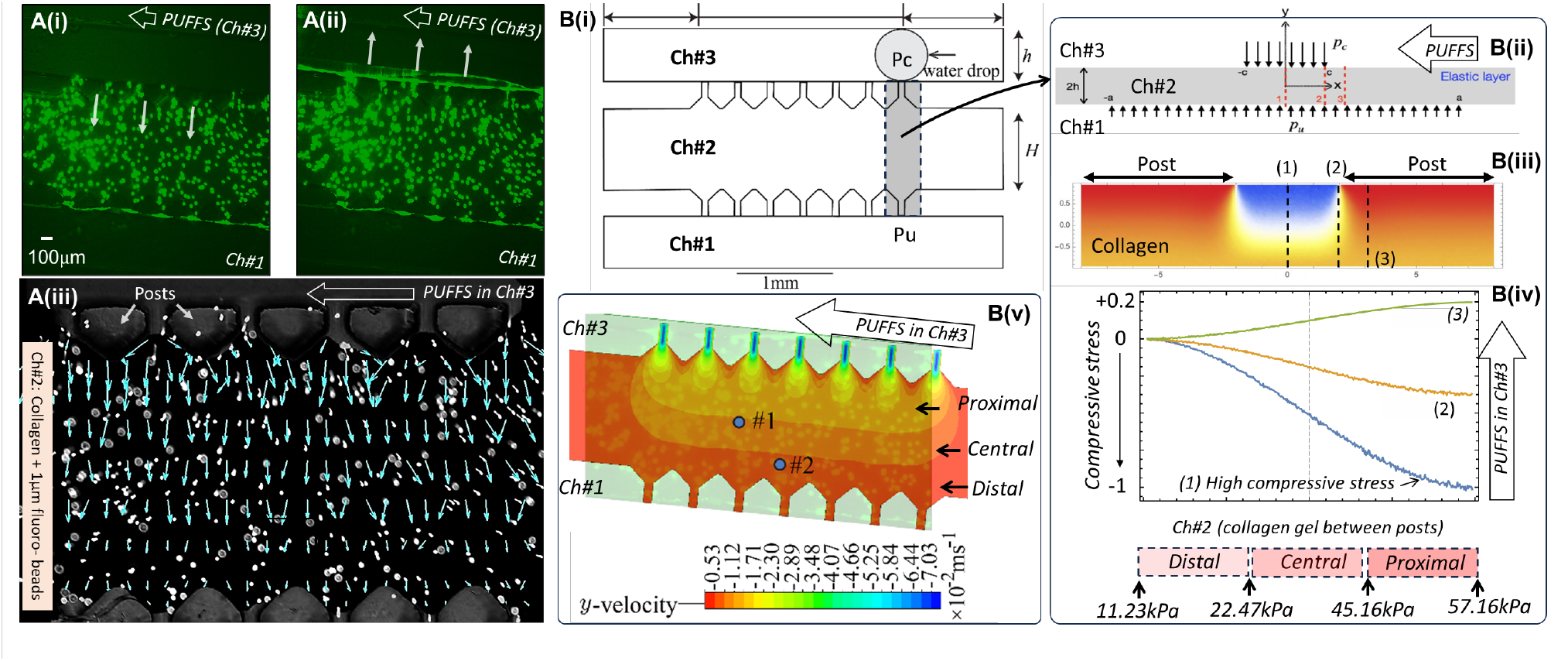
**A (i, ii)** Fluorescence images showing the movement of bead-laden collagen during PUFFS. **A (iii)** Vector map superimposed on brightfield image showing the magnitude of bead displacement. **(Bi)** Simplified geometry of the chip, **(Bii-iv)** Elastic modeling of collagen between two adjacent posts. **(Bv)** Velocity distribution within collagen during PUFFS. Scale bar=100 µm

### Viability, morphology and connectivity of 3D osteocyte networks subjected to PUFFS

Chips with MLO-Y4 osteocytes encapsulated within collagen were subjected to PUFFS, and their viability was assessed using a live dead staining assay. **(Fig.4A)** Results show a decrease in viability for both static (no PUFFS applied) and dynamic (PUFFS applied from Day 3-21, 0.33Hz) conditions. For instance, **Figure 4B** shows that viability decreased from 0.94±0.034 on Day 3 to 0.81±0.098 on Day 7 to 0.69±0.065 by Day 21 under static conditions. Under dynamic conditions, viability decreased from 0.93±0.016 on Day 3 to 0.88±0.018 on Day 7 to 0.81±0.086 by Day 21. At least 3 independent chips (samples) were used for this study and images were taken from the entire region in chamber #2.

**Figure 4.**
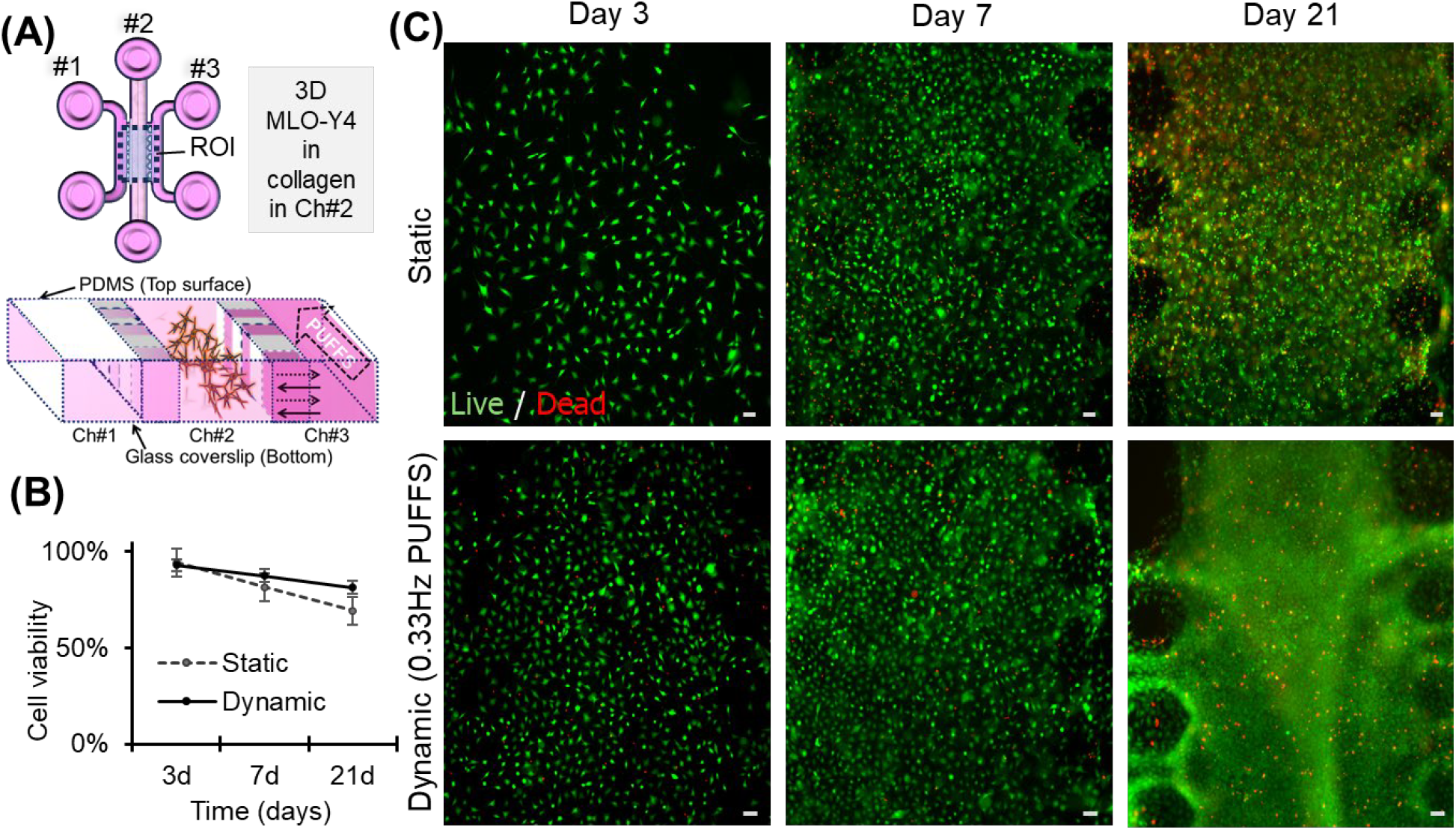
(A) Schematic of top view and cross-sectional view of the chip, (B) Cell viability as a function of culture duration under static and dynamic (PUFFS, 0.33Hz) conditions. (C) Representative fluorescence microscopy images captured from chamber 2 showing live (green) and dead (red) cells. Scale Bar=50 μm

MLO-Y4 morphology was assessed by staining cells for nucleus (blue) and f-actin (green). **(Fig.5A-D)** Since the total height of the collagen was ~250 µm, we took images at different z-depths from the bottom glass slide. Images taken from ~70 µm from the bottom were denoted by **(i)** in **Figs.5A-D**, and images taken from a z-plane close to the top PDMS surface (~200 µm from the bottom) were denoted by **(ii)** in **Figs.5A-D**. On Day 3, we observed many cells on the bottom, some cells in the middle section (i), but no cells in the top plane (ii). With longer culture durations, cells populated the entire depth of the collagen with highly spread-out cells in all planes on Day 21. This can be due to gravity-induced setting of cells during collagen gelation at early time-points, which eventually migrate into the collagen matrix by Day 7 and 21. Cell nuclei from captured images, from three independent chips, were used to calculate the number of cells per unit area. **(Fig.5D)** Results show an increase in cell number with culture duration under both static and dynamic conditions. Under static conditions, cell number increased from 139±92 (Day 3) to 591±102 (Day 7) to 2482±596 (Day 21), while under PUFFS conditions, cell number increased from 211±3 (Day3) to 587±352 (Day 7) to 2743±848 (Day 21). We found it challenging to identify connections between cells encapsulated within 3D collagen matrix, especially when cells are closely packed. Therefore, we use cell-nuclei separation distances as a criterion to generate 3D cell connectivity maps. Here, nuclei separation distances ≤ 50um were assumed to be connected cell-pairs (solid lines) while individual nuclei are represented as red-dots **(Fig.5E)**. The 3D maps provide a visual representation that shows an overall increase in cell number and cell migration into 3D collagen matrix, and cell-connectivity with longer culture durations. Maps show some cell aggregation in the dynamic condition as compared to the static condition which aligns well with observed cell morphology.

**Figure 5.**
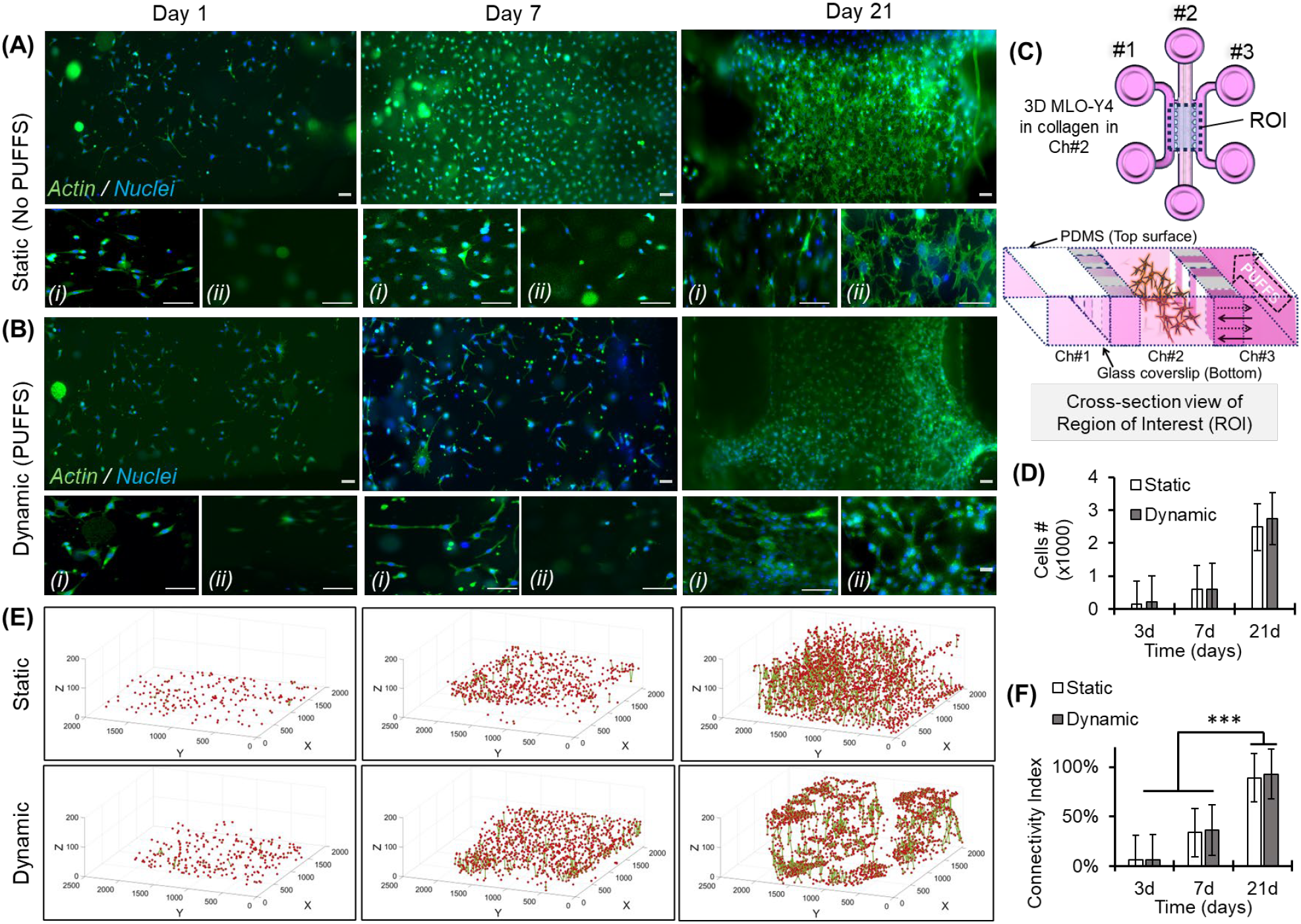
(A-B) Representative image of MLO-Y4 morphology in static and dynamic conditions; (i,ii) images taken at ~70µm and ~200µm from the bottom glass coverslip. (C) Schematic of chip, (D) Plot showing cell-number per unit area on Days 3, 7, and 21. (p<0.001). (E) 3D cell-connectivity maps; red dot = individual cell nuclei, green line=connection with adjacent cells. (F) Connectivity index on Days 3, 7 and 21. Scale bar=50 µm.

### Changes in gene expression in 3D MLO-Y4 networks under PUFFS (Figure. 6)

To assess the impact of PUFFS on gene expression by osteocytes, chips with 3D MLO-Y4 laden collagen gels were subjected to dynamic or static conditions for 7 or 21 days. Gene expression profiles of 10 mechanosensitive target genes (*Alpl, Gja1, Pdpn, Sost, Tnfsf11, Tnfsfr11b, Phex, Mepe, Dmp1* and *Fgf23*; **SI-6**) were analyzed via RT-qPCR, normalized to housekeeping genes (*Gapdh* and *Hsp90ab1*), and assessed via two-way ANOVA to assess statistical significance of differences between treatment groups as well as isolate the contributions of time (7d/21d), stimulus (Static/PUFFs), and the interaction of these terms. **(Fig.6)** Alkaline phosphatase (*Alpl*) is expressed by osteoblasts and early osteocytes and facilitates the deposition of mineralized matrix by hydrolyzing extracellular inorganic pyrophosphate, making it available for formation of calcium hydroxyapatite. *Alpl* exhibited significant variance from time (10.1%), stimulus (40.0%), and interaction (26.7%, all *p*<0.0001). Relative to 7d static controls, Alpl was downregulated in cells exposed to PUFFS for 7d (−3.73 fold, p=0.0049) with pronounced downregulation at 21d under both static (−26.3-fold, p<0.0001) and PUFFS (−6.48-fold, p<0.0001) conditions. There was no significant difference in Alpl expression in between cells exposed to PUFFS for 7d or 21d (−1.74-fold, p=0.1837). This aligns with ERK1/2-mediated suppression of Runx2-driven Alpl expression under shear stress,^62^ reflecting osteocyte maturation in vitro. Gap Junction Protein Alpha 1(*Gja1*) transcript encodes the protein connexin 43 (CX43), which is integral to osteocyte mechanotransductive function in that it enables transmission of small signaling molecules (e.g. Ca^2+^) between interconnected osteocytes.^63–64^ Gap Junction Protein Alpha 1 (Gja1/Cx43) showed no significant differences, suggesting baseline gap junction integrity is transcriptionally stable, as indicated by immunofluorescence. Podoplanin (*Pdpn*), also known as protein E11, is implicated in the initial stages of osteocyte differentiation and is essential for the formation of dendritic processes and has been suggested to act as a sensor for bone damage.^65–66^ Expression of *Pdpn* was suppressed by PUFFS (−3.06-fold, p=0.0070 at 7d; −5.52-fold at 21d, p<0.0010). Relative to 7d static controls, downregulated Pdpn (−10.69-fold, p<0.0001) at 21d. There was no significant difference in *Pdpn* expression between cultures exposed to PUFFS for 7d and 21d (p=0.1879). These changes appeared to be driven by stimulus (50.1%) and interaction of time and stimulus terms (21.3%, p<0.0001). This contrasts with research demonstrating that *Pdpn* increases with in vivo loading, enhances osteocyte connectivity in dendritogenesis. Reduced *Pdpn* as observed here may indicate excessive shear stress activating RhoA/ROCK pathways that trigger cytoskeletal retraction of dendritic processes.^67^ Alternatively, this reduction of *Pdpn* with PUFFS may indicate a matrix-driven feedback mechanism because collagen gel stiffness may insufficient to support dendrite extension, despite mechanical stimulation.^68^ Unloading of bone promotes the secretion of Sclerostin, encoded by the *Sost* gene, by osteocytes acting as a negative regulator of bone formation by inhibiting the Wnt/β-catenin signaling pathway.^69–71^ Expression of (*Sost*) was predominantly influenced by the interaction of time and stimulus variables (89.05%, *p*<0.0001). In comparisons to 7d Static controls, exposure to PUFFS for 7d for did not significantly affect *Sost* expression (1.19-fold p=0.9994) but was downregulated relative to 7d in for both static (−14.94-fold, p<0.0001) and PUFFs (−13.23-fold, p<0.0001) conditions. In vivo loading suppresses *Sost* via Piezo1 activation,^72^ it is possible that the less stiff environment provided by the collagen gel limits prevented mechanosensitive Piezo1 activation, leaving time-dependent silencing dominant. While Sost downregulation should activate Wnt signaling, the absence of anabolic gene upregulation (e.g., Alpl, Dmp1) suggests that reduction of Sost may be compensated by Secreted Wnt antagonists (e.g., Dkk1) not assayed here,^68, 73^ or via mechanically activated signaling through the Wnt/Ca^2+^ or Wnt/PCP pathways.^74^ *Fibroblastic Growth factor 23* (*Fgf23*) is expressed by osteocytes at the most advanced stage of differentiation. Mechanical strain has been suggested to modulate expression of *FGF23*, which acts on the kidney to regulate systemic phosphate and vitamin D metabolism.^75–77^. While *Fgf23* was detected under both static and PUFFs conditions after 7d in the device, expression was not significantly different between treatments (+1.48-fold, p=0.9998). In contrast, *Fgf23* was barely detectable in MLO-Y4 cultured for 21d under either static or PUFFs conditions. Fibroblast Growth Factor 23 (Fgf23) became undetectable at 21d, correlating with Phex downregulation (−7.77-fold static 21d). Mechanical loading *in vivo* requires Phex-mediated cleavage of MEPE for Fgf23 maintenance. Mineral-free collagen gels, as studied here may impair feedback via the Phex-Fgf23-MEPE axis mimicking osteocyte dedifferentiation and disrupted phosphate homeostasis.^78^ Receptor activator of nuclear factor κB ligand (*Rank-L* or *Tnfsf11*) and osteoprotegerin (*Opg* or *Tnfsfr11b*) are secreted by osteocytes to modulate osteoclast development and resorptive activity. While *Rank-L* promotes osteoclast differentiation and bone resorption, *Opg* acts as a decoy receptor that neutralizes *Rank-L* to fine-tune the osteoclastic component of mechanoadaptation.^79–80^ Variance of *Rank-L* expression stemmed from time (16.4%), stimulus (17.3%), and interaction (39.5%), with PUFFS reducing *Rank-L* at 7d (−5.58-fold, *p*=0.0052). Mechanical suppression of *Rank-L* mirrors *in vivo* loading’s anti-resorptive effects via *Mepe* upregulation.^81^

**Figure 6.**
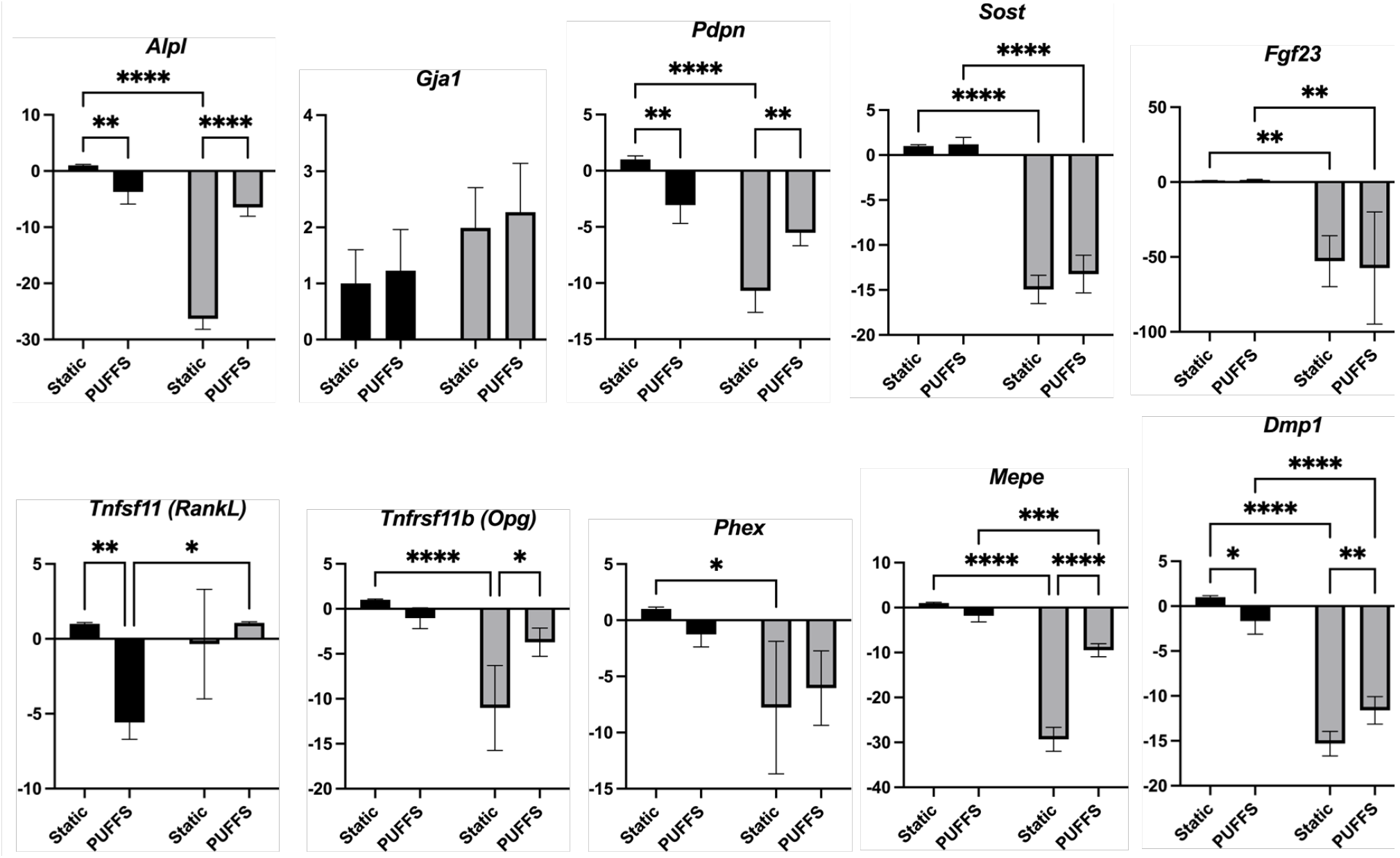
Gene expression by RT-qPCR for 10 osteocyte genes. Data shown are mean +/-SD of n=3-5 replicates, and brackets show statistically significant differences (p<0.05) in gene expression between treatment groups by 2-way ANOVA (Time^*^treatment).

Osteoprotegerin (Opg/Tnfsfr11b) increased 2.97-fold at 21d PUFFS (*p*=0.0164), driven by stimulus (40.9%, p=0.0009) and interaction (16.6%, p=0.0077). Sustained OPG elevation aligns with mechanical promotion of decoy receptor production to buffer *Rank-L*.^82^ Prolonged mechanical stimulation enhances OPG production, consistent with *in vivo* loading suppressing resorption.^81^ Pulsatile fluid flow in MLO-Y4 cells has been shown to increases MEPE, lowering RANKL/OPG ratios and inhibiting osteoclastogenesis.^81^ Taken together these results suggest that prolonged mechanical stimulation enhances OPG production, consistent with *in vivo* loading suppressing resorption. Phosphate-regulating neutral endopeptidase (*Phex*) gene encodes a zinc metalloendopeptidase expressed in osteocytes that is involved in bone mineralization and phosphate homeostasis via its influence on *FGF23* expression. ^83–84^ In two-way analysis, only treatment contributed significantly to the observed variance (44.1%, p=0.0040). *Phex* showed stimulus-driven downregulation (−7.77-fold static 21d, *p*=0.0110), implicating mineral-free collagen gels in disrupting the Phex-Fgf23 axis critical for phosphate regulation.^78^ Osteocytic expression of Matrix extracellular phosphoglycoprotein (*Mepe*) is modulated by mechanical stress, suggesting its involvement on adaptation to mechanical loading. Time (10.1%), stimulus (50.2%), and interaction (17.9%) significant. PUFFS reduced *Mepe* at 21d (−7.03-fold vs. static). In pairwise comparisons, *Mepe* was suppressed by PUFFS at 21d (−7.03-fold, *p*<0.0001), *Mepe* expression was also significantly reduced between 7d and 21d timepoints in both static (−29.3fold, p<0.0001) and PUFFS treated cultures (−7.03 fold, p=0.0006) in PUFFs treated cultures. This pattern contrasts to the induction of *Mepe* observed with mechanical loading in vivo. This paradox may reflect overload stress or absent mineral feedback. Similar to *Mepe*, expression of the *Dentin matrix protein 1 (Dmp1)* transcript by osteocytes is upregulated in response to mechanical loading, though the Dmp1 protein is involved in both positive and negative regulation of matrix mineralization, and dependent upon post-translational modification and cleavage into fragments of varying function.^84–86^ Furtheremore both *MEPE* and *DMP1* proteins are substrates of *PHEX*, whose proteolytic activity releases acidic serine aspartate-rich *MEPE*-associated motif (ASARM) peptides that bind hydroxyapatite and negatively regulate further matrix mineralization.^87–88^ Two-way analysis of data for *Dmp1* expression showed that while time did not contribute significantly to the observed variance (0.1%, p=0.3980), stimulus (77.8%, p<0.0001) and the interaction of time and stimulus (4.6%, p=0.0002 were significant factors. There was no significant difference in *Mepe* expression between static and PUFFs treated cells at 7d (−1.83-fold, p=0.1690), at 21d PUFFs significantly reduced expression (p<0.0001) compared to 21d static treatment. Dmp1 expression was reduced between 7 and 21d in culture for both static (−15.31-fold, p<0.0001) and PUFFS conditions(+3.09-fold, p<0.0001). Dmp1 was downregulated by PUFFS (−5.17-fold at 21d, *p*=0.0039), opposing in vivo loading’s enhancer-driven Dmp1 activation,^78^ suggesting non-physiological PUFFS parameters.

The application of dynamic mechanical stimulation (PUFFS) to MLO-Y4 osteocyte-like cells in a 3D collagen bioreactor system revealed complex temporal and stimulus-dependent regulation of genes governing bone mineralization, osteocyte differentiation, and osteoclast-osteoblast coupling. Key patterns include: (1) Time-driven suppression of mineralization regulators (*Alpl, Mepe, Dmp1*) independent of mechanical stimulus^82^ (2) PUFFS-mediated anti-catabolic effects via RANKL/OPG modulation, (3) Culture duration dominance over *Sost* expression, and (4) Loss of mature osteocyte markers (*Fgf23*) at later timepoints. Despite *Sost* suppression, anabolic genes (*Alpl, Dmp1*) remain low. This mirrors β-catenin-independent Wnt signaling (e.g., Wnt/Ca^2+^) activated by mechanical stress, bypassing transcriptional targets like *Alpl*.^62^ Culture duration eclipses mechanical effects on Sost and Fgf23, highlighting limitations of prolonged in vitro osteocyte models. Mechanistically, PUFFS aligned with ERK1/2-mediated Runx2 suppression (Alpl) and RhoA/ROCK-driven dendrite retraction (*Pdpn)* but diverged in Wnt (*Sost*) and mineralization pathways, perhaps due to collagen gel constraints. These findings underscore the need to optimize mechanical parameters (e.g., shear stress magnitude: 0.5–3 Pa; ^72^ and incorporate mineral phases to better recapitulate *the in vivo microenvironment of osteocytes*.

### Expression of key markers in 3D MLO-Y4 osteocyte networks

We stained for key proteins related to osteocyte biology. **(Fig.7)** Sclerostin (Sost), widely used to identify osteocytes, was stained for chips in both static and dynamic conditions on Days 7 and 21. Next, we stained for αvβ3 integrin, a receptor on osteocytes that facilitates attachment to collagen, and gap-junction protein Cx43 that is known to facilitate mechanical stimuli-evoked calcium ion signaling. For both Cx43 and αvβ3, we observed that the staining was distributed over the entire cell surface and higher levels of staining can be seen for the dynamic group as compared to static control. Control experiments for all immunostains (exlusion of primary antibodies) show little-to-no non-specific fluorescence signals **(SI-7)**

**Figure 7.**
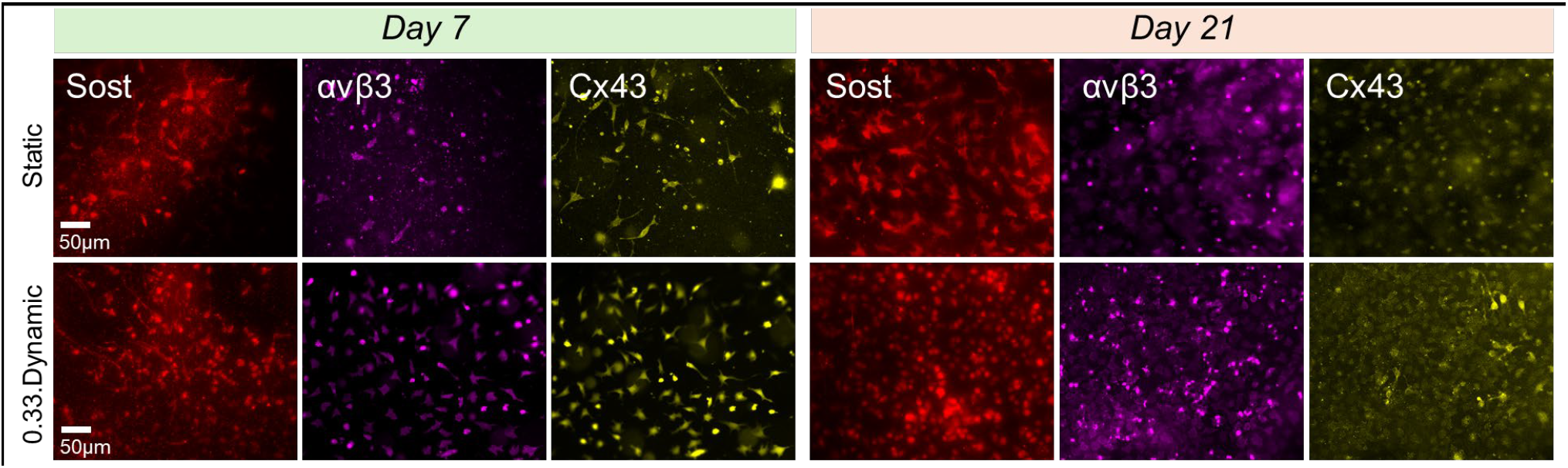
Representative fluorescent images showing the expression of sclerostin (red), connexin-43 (Cx43 - yellow) and αvβ3 (pink) under static and dynamic conditions at day 7 and day 21. Scale bar = 50µm).

### Real-time monitoring of calcium signaling within 3D osteocyte network under PUFFS

Fluo-4AM calcium staining in combination with time-lapse fluorescence microscopy was used to capture the changes in calcium intensity of MLO-Y4-laden collagen in chamber #2. For every experiment, the baseline signal was captured under no stimulation conditions for ~40s followed by the application of PUFFS (0.33 Hz) for ~60seconds in chamber #3. For every experiment, the average magnitude of baseline signals (0-40s) was used to normalize the calcium signals. Representative fluorescence images and video captures of MLO-Y4 calcium signaling in chamber #2 show the propagation of calcium signal across the MLO-Y4 network, starting from chamber #3 to chamber #1. **(Fig.8A, Movie S2)** We identified individual cells within the network and plotted changes in calcium intensity as a function of time. **(Fig.8B-C)** We observed similar signals for both static and dynamic conditions. All MLO-Y4s showing oscillatory response show one of two signaling profiles (**Fig8C**). One profile type returns to baseline fluorescence values after every PUFFS-induced calcium spike while the other type exhibits a gradual increase in the overall fluorescence and does not come back to baseline signals even after PUFFS is stopped. We used *PeakFinder* (MATLAB) to assess the oscillatory frequency of cell-laden MLO-Y4s in chamber 2. **(Fig.8D)** We found that the frequency increase from 0.39±0.83 Hz (Day 7) to 0.46±0.22 Hz (Day 21) under static conditions. On the other hand, for chips subjected to daily PUFFS, the oscillation frequency decreases from 0.45±0.04 Hz (Day 7) to 0.43±0.33 (Day 21). We further characterized the number of MLO-Y4 showing oscillatory signals. **(Fig.8E)** We found that, for static culture, MLO-Y4s exhibiting oscillatory response increased from ~46% (Day 3) to ~99% (Day 21) while for chips subjected to daily PUFFS (0.33 Hz), cells exhibiting oscillatory response increased from ~70% (Day 3) to ~83% (Day 21). Since both the mechanical deformation of the collagen matrix and cell-to-cell gap-junction based signaling could modulate their response, we repeated this experiment in the presence of a gap junction Cx43 blocker (GAP26). Briefly, on Days 7 and 21, the GAP26 solution is pipetted in side-chambers for 45 minutes before applying PUFFS (0.33 Hz, 60s) in chamber #3. Results show that, for static conditions, the total number of MLO-Y4s that exhibit an oscillatory response decreased from 37% (Day 7) to 21% (Day 21) while for chips subjected to daily PUFFS (0.33Hz), oscillatory cells decreased from 28% (Day 7) to 16% (Day 21). After blocking with GAP26 (Cx43 inhibitor), we saw an overall decrease in the number of cells showing the oscillatory response. This indicates that at early time-points, when connectivity is low, inhibition of gap junction does not play a significant role; this could mean that the responses we observe are more due to mechanical deformation. At longer culture durations (Day 21), with more cell-to-cell connectivity, there is a significant drop in the number of cells showing an oscillatory response, which indicates a significant role of gap-junction based signaling in addition to matrix deformation induced signaling responses recorded during PUFFS.

**Figure 8.**
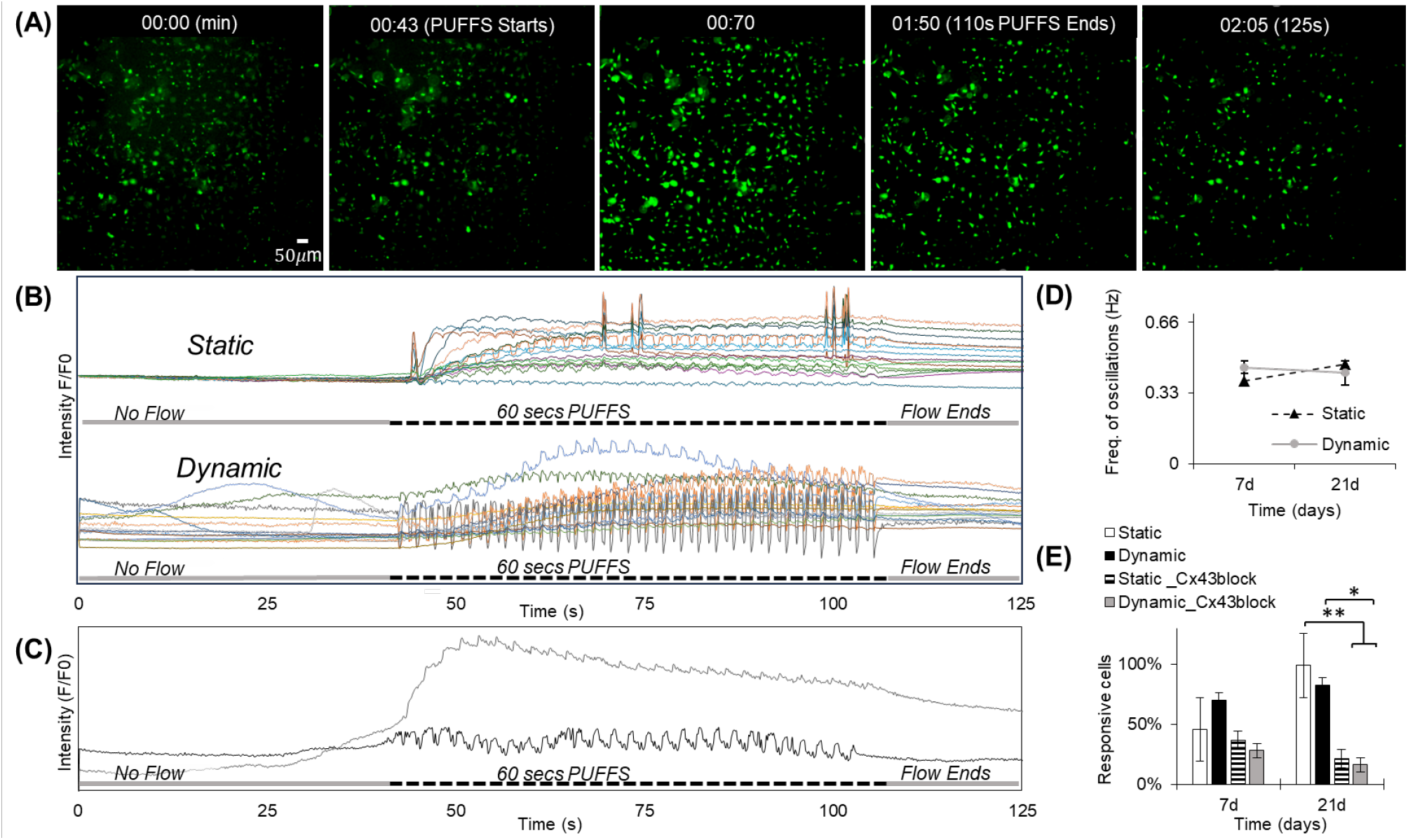
(A) Snapshots of fluorescence images showing propagation of calcium signaling within MLO-Y4-laden collagen in chamber 2. (B) Representative normalized calcium signaling profiles before, during and after the application of PUFFS, (C) Plots showing typical single cell responses. (D) Plot showing frequency of signal oscillations for static and dynamic chips, (E) Number of responsive cells with oscillating signals in the presence and absence of Cx43 gap junction inhibitor for static and dynamic chips for Days 7 and 21.

### Characterization of calcium signaling within sub-regions of MLO-Y4-laden collagen

We investigated if regions closer and farther away from PUFFS elicit similar signaling responses. To do so, chamber 2 was divided into three regions: proximal (0 - 285 µm), central (286-570 µm), and distal (571-855 µm) **(Fig.9A)**. First, the number of cells exhibiting oscillatory response were characterized using the *PeakFinder*.*m* (MATLAB) as explained in the methods section **(Fig.9B)**. For Day 3, in proximal subregion, responsive cells were 21±0.07% (static) and 28±0.08% (dynamic), while central subregion showed 24±0.16% (static) and 21±0.13% (dynamic) of responsive cells, while for distal subregion, 12±0.14% (static) and 12±0.06% (dynamic) were recorded. For Day 7, in proximal subregion, responsive cells were 14±0.12% (static) and 19±0.02% (dynamic), while central subregion showed 16±0.15% (static) and 25±0.11% (dynamic) of responsive cells, while for distal subregion, 16±0.05% (static) and 27±0.15% (dynamic) were recorded. For Day 21, in proximal subregion, responsive cells were 29±0.05% (static) and 30±0.16% (dynamic), while central subregion showed 31±0.09% (static) and 32±0.13% (dynamic) of responsive cells, while for distal subregion, 40±0.05% (static) and 21±0.04% (dynamic) were recorded. To assess how the signaling properties adapt to PUFFS, we applied multiple rounds of Fluo-4AM calcium dye staining on the same sample, however, this resulted in significant cell death. As a result, we tried to compare calcium signaling characteristics from independent chips (samples) using calcium dye staining as an end-point assay. Thus, chips were analyzed for each time-point (Days, 3, 7, and 21), and their signaling properties were compared. First, single cells in each sub-region exhibiting oscillatory calcium signal during PUFFS were pooled together to obtain a cumulative signal that could represent the proximal, central and distal sub-regions. To extract a representative calcium signal from each of these sub-regions, xcorr function (‘signal/SignalSimilaritiesExample’) (MATLAB) was used to cross-correlate calcium signals from all single-cells located within each sub-region. The xcorr function measures the similarity between two signals at a specified time length and computes the lag differences, where zero lag indicates matching signals. The Excel SORT function was then used to reorder responses based on distance from PUFFS and group cells with similar responses together. Only cells exhibiting oscillatory response were analyzed with 80-400 cells per region obtained from 3 independent chips. Oscillatory cells from proximal, central, and distal groups were averaged into a single response; thus, each chip had 3 regional responses. **(Fig.9C)** For each regional response, the start and stop of PUFFS are indicated by arrowheads. Despite maintaining consistency in threshold settings for all chips and analyzing many single-cell responses (80-400), we found that calcium signals show large variations in regional responses making direct comparison between chips challenging. We also conducted this experiment in the presence of GAP26 (Cx43 inhibitor) to check if signaling properties are affected, but due to large signal variations between chips, a direct comparison was not possible.

**Figure 9.**
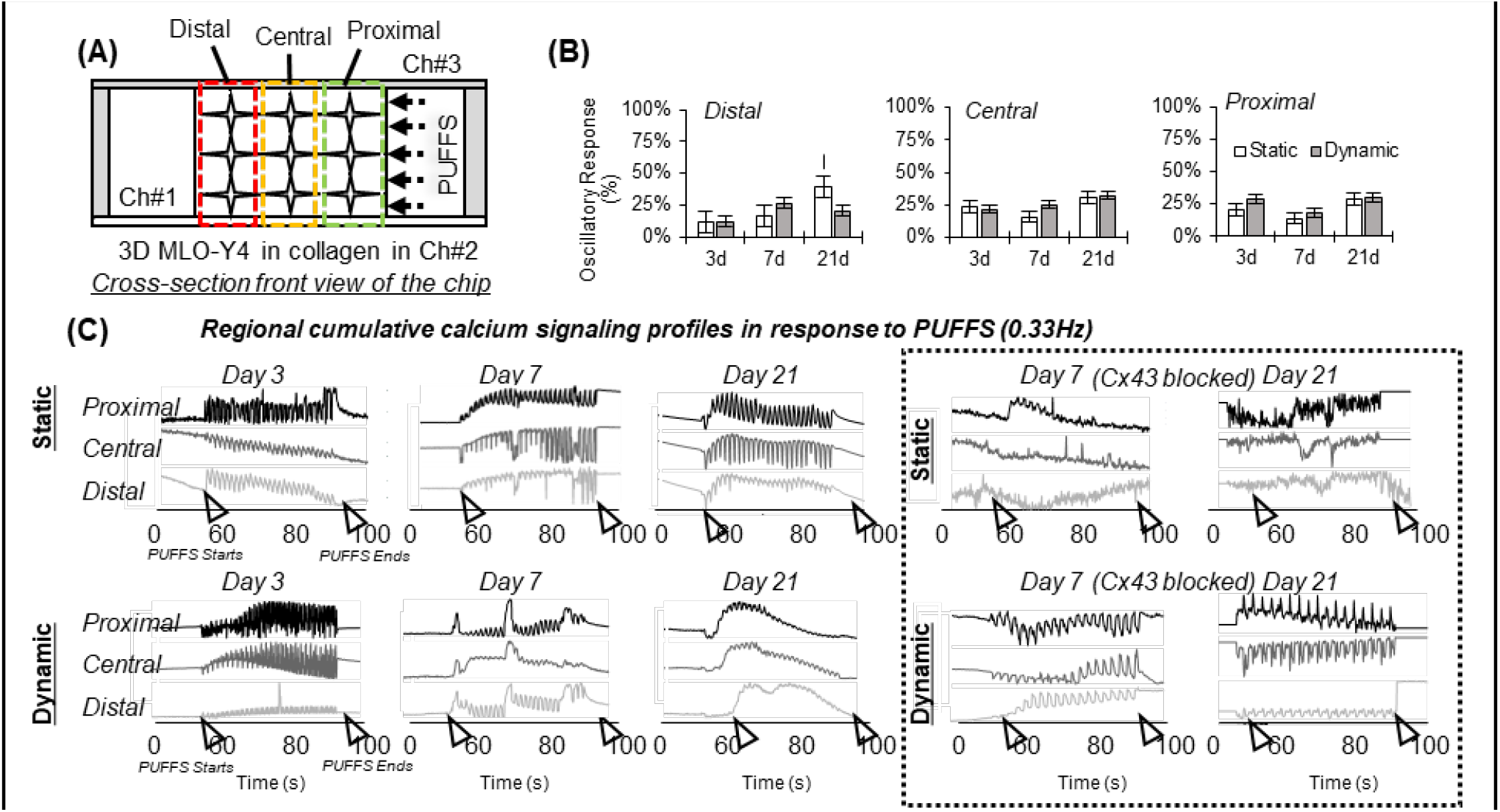
(A) Schematic showing proximal, central, and distal sub-regions with respect to PUFFS. (B) Plots showing number of MLO-Y4s with oscillatory responses within each sub-region for Days 3, 7 and 21. (C) Left: Representative signaling profiles from various sub-regions before, during, and after PUFFS. Right: Representative plots when gap junction was blocked on Day 7. Each signal is captured from an independent chip with application of PUFFS for 60s (marked by black arrowheads).

In another study, we tested whether sequential application of PUFFS at 0.33Hz and 1.66Hz is possible and whether a stable collagen barrier is retained**(Fig. 10A, B)**. Process workflow shows the application of PUFFS to both static control and daily PUFFS groups first at 0.33 Hz, followed by a 15-minute rest period before a second application of PUFFS at 1.66 Hz. Results show that osteocytes show signaling spike frequency are lower than the PUFFS frequency of 1.66 Hz. (**Fig.10C)** Day 7 static conditions had 55±0.37% response cells and 61±0.03 for PUFFS (1.66 Hz) which increased to 75±0.41% (static) and 79±0.22% (dynamic) by Day 21. We also carried out identical experiments using Cx43 gap junction blocker (**Fig.10D**). Unlike the first PUFFS application at 0.33 Hz, blocking by GAP26 did not greatly reduce the oscillatory responses for dynamic conditions during sequential PUFFS application at 1.66 Hz.

**Figure 10.**
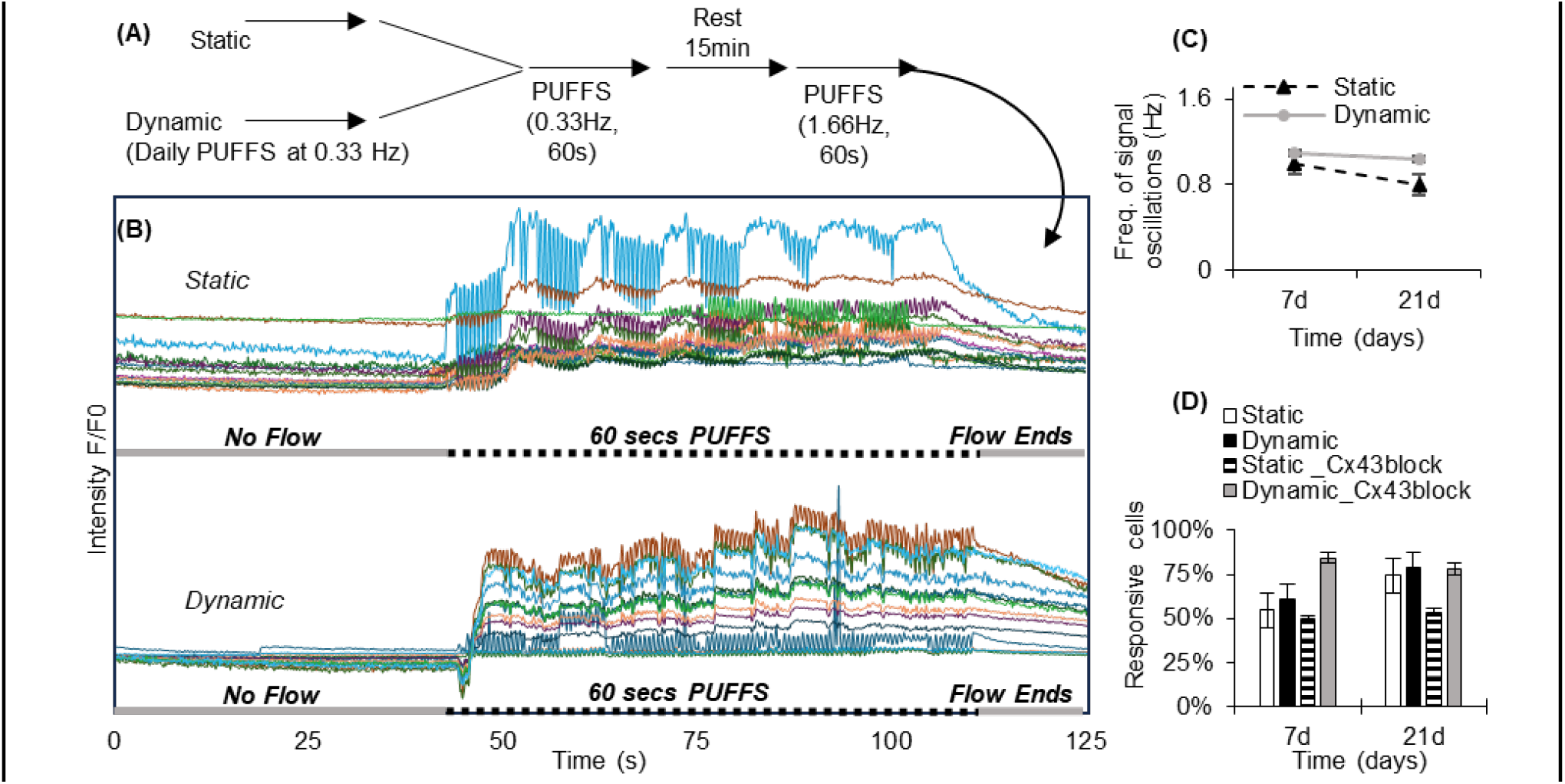
(A) Sequential application of PUFFS at 0.33 Hz (60s), then rest for 15-minutes followed by PUFFS at 1.66 Hz (60s). (B) Representative normalized calcium signaling profiles before, during and after the application of PUFFS, (C) Plot showing frequency of signal oscillations for static and dynamic chips, (E) Number of responsive cells, with oscillating signals, in the presence and absence of Cx43 gap junction inhibitor for static and dynamic chips for Days 7 and 21. (PUFFS, 1.66 Hz)

In this work, we designed and developed a new in vitro model that can be used to study dynamic signaling within 3D osteocyte network when subjected to defined pulsatile mechanical stimuli. Due to the difficulties in obtaining and culturing primary human osteocytes, we choose to work with immortalized murine osteocyte MLO-Y4, a cell-line widely used in the field due to its dendritic morphology, sensitivity to fluid-flow and biochemical stimuli, proven utility in previous publications.^30–37, 89–90^ Since the organic portion of bone tissue ECM is mostly collagen (90%), we choose to make 3D osteocyte networks using collagen type I and chose stimuli-evoked calcium (Ca^2+^) transients^91–95^ as a proxy for real-time signaling. We choose PUFFS stimuli at 0.33 Hz and 1.66Hz (15 mins/day) based on previous work that found bone cells respond favorably to repeated short bursts of flow after 10 to 15 minute rest periods.^96–97^ A concentration of 2.5 mg/ml was chosen for this study, as it allowed long-term culture of MLO-Y4s under both static and dynamic conditions. Day 21 was chosen as the endpoint, as collagen starts to become unstable beyond this time-point, due to detachment from the side walls, especially under PUFFS (1.66 Hz). Studies have recorded a variety of responses of osteocytes to fluid flow stimuli.^98–100^ In our studies, when PUFFS was applied, both mechanical deformation of collagen and gap-junction based signaling contribute to calcium signaling profiles. Moreover, the percentage of MLO-Y4s that respond to PUFFS in the form of oscillatory responses (responsive cells) vary with static and dynamic samples and increased cell number and connectivity by Day 21. In this work, PUFFS-evoked signaling were short-term experiments lasting less than 30mins after adding commercial Fluo-4AM calcium dye followed by a semi-automated thresholding method to analyze signaling profiles of 100s of individual MLO-Y4s. However, due to large variations in calcium signaling profiles, we found it challenging to directly compare results across different chip samples. To address this challenge, we performed Fluo-4AM staining on the same chip every other day, however we did not pursue this further due to decrease in cell viability. In the future, stably transfected variants of MLO-Y4s that express fluorescent fusion proteins labeling the plasma membrane and a genetically encoded calcium indicator could facilitate longitudinal calcium signaling using the same chip. Due to the large amount of real-time signaling data generated, automated mapping of single cells along with functional connectivity graph, pairwise co-activity (correlations), and co-occurrences (signal synchrony) should be used in the future. The numerical simulation used in this work also has some limitations. The simulations used a planar representation of the experimental design, which does not fully capture the three-dimensional nature of the actual system. Additionally, the material properties used in the simulation, such as the viscosity and density of the collagen gel, were based on literature reviews and may not accurately reflect realistic variations in collagen. Despite these limitations, our work designed and developed a new chip to enable the study changes in 3D osteocyte networks subjected to PUFFS. We envision that this minimally invasive chip can be potentially extended to other cell types, and could be used to test a range of biophysical and biochemical stimuli.

## MATERIALS & METHODS

### Design and fabrication of multi-chambered microfluidic PDMS chips using 3D printed molds

Molds were printed using a prepolymer solution consisting of 40 mL of poly(ethylene glycol) diacrylate (PEGDA, average M_n_ 250) and 0.25% of photoinitiation agent (Irgacure 819, Sigma-Aldrich), 0.01% of TEMPO (Sigma-Aldrich) and 0.5% of 2-Isopropylthioxanthone (ITX, Tokyo Chemical Industry). The final prepolymer solution was protected from light with aluminum foil and thoroughly mixed using a vortex for 10 minutes. Glass slides (25 x 75 x 1 mm, Fisher brand) were pre-cleaned with piranha solution (H_2_SO_4_ and H_2_O_2_; 7:3, stirred for 30 minutes at 125 rev/min), washed with ethanol and water until reaching a neutral pH, and dried in vacuum oven at 65°C. The glass slide surface was further modified and stirred (125 rev/min at 50°C) in 3-(trimethoxysilyl)propyl methacrylate (TMSPMA; Sigma-Aldrich) and Toluene (Sigma-Aldrich) (9:1), then dried in a vacuum at 65°C. Modified glass slide was sliced using carbide-metal etching pen into 4 pieces, and each glass piece was adhered to an aluminum print block using a double-sided tape and screwed into the printer stage head. CAD files of the PEGDA master molds were generated using Fusion 360 and exported as stl files, then imported into MATLAB to develop sliced image files for the master mold and exported as PNG files (i.e., black and white resolution 1080 x 1080). Image slices were uploaded as virtual masks into the digital micromirror device (DMD) software controlled by the LabView code. For 3D printing we utilized the Digital Light Projection (DLP, development kit 1080p 9500 UV, Texas Instruments, USA) platform designed, and custom built in Soman group.^90, 101^ Optical settings consisted of a rotating diffuser to minimize light speckles, a z-stage (25 mm Compact Motorized Translation Stage, ThorLabs), and ultraviolet light (400 nm) laser source (iBEAM SMART 405, Toptica Photonics). This setup was used to print 250 µm thick PEGDA master molds onto the treated glass slides. The height of molds was measured with a digital caliper (Mitutoya). Multiple master molds were printed with minimal batch variation and were used to make chips using replica casting. Polydimethylsiloxane (PDMS) elastomer was mixed with curing agent (Sylgard 184, Dow Corning Silicone Elastomer) at 10:1 for 10 minutes and degassed in a desiccator to remove air bubbles. The PDMS precursor was cast on at least 5 PEGDA master molds evenly spread in a 100 x 15 petri dish and kept in a vacuum oven at 70°C overnight with vacuum shut off to avoid bubble accumulation. After cooling, the PDMS reverse molds were peeled from the PEGDA master molds. A 2 mm biopsy punch was used to create 6 holes for the inlet/outlet ports. The edges of the PDMS molds were cut and bonded to glass coverslips (22 mm x 22 mm x 0.17mm, PCS-1.5-2222, Mattek), pre-cleaned using an overnight acid wash (30% Hydrochloric acid). Prior to bonding, PDMS molds and glass coverslips were plasma treated (PE-50 model, Plasma Etch Inc.) and heated on hotplate at 150°C for an hour. Before use, the chips were sterilized by incubating in 100% ethanol followed overnight exposure of UV radiation in a BSL–2 cell culture hood.

### 3D osteocyte culture within three-chambered microfluidic PDMS chips

Before incorporating MLO-Y4 cells in the chips, chips were surface-modified using an established protocol. Briefly, 2 mg/ml polydopamine (PD) (Sigma-Aldrich, H8502) solution was pipetted in the chamber 2 and incubated at room temperature for 24 hours, then washed with PBS (3x). Then chip surfaces were incubated in 0.01% poly-l-lysine solution (PL) (Sigma-Aldrich, P4707) for 15 minutes at room temperature and washed three times with PBS followed by another coating of 0.15 mg/ml rat tail type 1 collagen (RC) for an hour at room temperature. The central channel was washed three times with PBS, dried, and sterilized under UV radiation for 45 minutes before incorporating cell solution in the chip. MLO-Y4 osteocyte cell line (Kerafast, Inc. Boston, MA), maintained in alpha-MEM (12571063, Gibco), containing L-glutamine, 1% penicillin/streptomycin, 2.5% fetal bovine serum, and 2.5% calf serum, was cultured in flasks coated with 0.15 mg/ml rat tail type 1 collagen (Advanced Biomatrix) at 37^°^C under a humidified atmosphere of 5% CO_2_. Upon reaching 75-90% confluency, cells were trypsinized (0.25%) and resuspended in media and mixed with collagen solution. To prepare the collagen solution, 1333 µL of bovine type 1 collagen (5225 bovine, Advanced BioMatrix, 6 mg/ml) was mixed with 414 µL 10x HBSS (#14065056, ThermoFisher), and 236 µL neutralizing agent (Advanced Biomatrix). For all chips, 10 µL of cell solution (~18,000 cells per chip) was pipetted in the central chamber (Ch#2) and incubated at 37°C for 30 minutes to achieve gelation of collagen with encapsulated MLO-Y4s. Media was pipetted in chambers 1 and 3 (side-chambers on either sides of collagen barrier) and replenished daily for Days 3, 7 or 21.

### Rheological characterization of collagen gels (SI-2)

TA Instruments Discovery Hybrid Rheometer DHR-3 equipped with a lower Peltier plate and 20mm crosshatched upper and lower geometries (TA Instruments, New Castle, DE, USA) was used to assess the mechanical properties of collagen gels. Gel slabs of approximately 1mm thickness were prepared by thermally crosslinking a 2.5 mg/mL bovine collagen solution at 37°C for 30 minutes. Prior to analysis, excess superficial water was removed from the gels via gentle blotting with a Kimwipe. Gel slabs were then placed on the rheometer and manually trimmed to yield 20 mm diameter discs. Samples were allowed to equilibrate at 37°C for 180 seconds prior to a frequency sweep experiment from 0.1 to 100 Hz with 2% strain. Storage and loss moduli are reported as the mean values measured at 1 Hz. All rheological measurements were performed in triplicate.

### Design and optimization of PUFFS setup

The overall setup involves two RP peristaltic pumps, (#RP-HX01S-1H-DC3VS, Takasago for PUFFS at 0.33Hz, and #RP-QX1.5S-1H-DC3V, Takasago for PUFFS at 1.66Hz), flow controller, and tubing that connect to the inlet and outlets of a chip. During flow stimulation experiments, the settings for PUFFS were adjusted with a controller and monitored at the motor’s frequency reading near 1200 ± 30 Hz (or Pulse Per Second (PPS) by manufacturer). Based on the manufacturer’s reduction gear ratio (1/50) and 0.015 conversion factors, the actual fluid frequency approximated to 0.33 Hz. **(SI-8)** For the other pump, similar adjustment was performed to achieve PUFFS at 1.66Hz. Pressure measurements were performed for 30 minutes with a relative and differential pressure transmitter (Type 652, Huba control, pressure range: 0-100 kPa). We confirmed 30 kPa as the approximate generated pressure from Takasago’s peristaltic pumps based on pressure measurements collected within 15 minutes. The pump positive flow tubing was connected to the P1 higher pressure (lower port) of the transmitter. The tubing that administered negative flow (suction) pulled DI Water from a petri dish. We used a G⅛ male to ¼” barb fitting at the P1 port followed by 50 mm of ¼” tubing and converted to 0.8 mm OD barb fitting using a ¼” barb to ¼-28 NPT fitting, a ¼-28 NPT union and a ¼-28 NPT to 0.8 mm OD barb. The final 0.8 mm OD barb fitting was attached to the peristaltic pump outlet through 90 mm of 1 mm ID tubing. The pump inlet tubing was submerged in DI water within a petri dish. Voltage output from the pressure transmitter was monitored with PD603 Low-Cost OEM Process Meter (Sabre Series). The tubing was filled with DI water so there was no air in the lines and the pressure transmitter was left to equalize until it read 0V while the pump was off. Once zeroed, the pump was turned on and proceeded for 15 minutes to collect continuous readings. To estimate how long PUFFS takes to enter and exit the channel, we intentionally allowed air to enter the tubing during PUFFS. Using air-bubble, the PUFFS velocity inside chamber 3 was calculated as ~0.011 m/s. Before applying PUFFS to chamber 3 of the chips, peristaltic pumps were pre-sterilized by perfusing 70% ethanol for 5 minutes and dried for 15 minutes. On Day 0, MLO-Y4-laden collagen was crosslinked in chamber 2 of the chips. On Day 1, the media was replenished, and PUFFS was applied for 15 minutes daily until target endpoint was reached (Day 3, 7 or 21). All perfusion experiments were performed at room temperature in a sterilized biosafety cabinet level 2 (BSL-2). After PUFFS, the media was replenished and chips were cultured under standard conditions (37^°^C, 5% CO_2_).

### Recording of calcium signaling and analysis

PUFFS evoked calcium signaling within MLO-Y4-laden collagen were tested on Day 3, 7 and 21. For each timepoint, 3 independent chips were tested. Device stabilizer was generated via Fusion 360, exported as a stl. file and 3D printed by fused deposition modeling (FDM, Bambu Lab P1P equipped with smooth PEI plate) using poly(lactic acid). **(SI-1)** During printing, the temperature for nozzle (0.4 mm) and bed plate was adjusted to 250°C and 65°C respectively. During testing, chips combined with 3D printed stabilizer were placed in a 35 x 10 mm petri dish, and calcium dye solution (500 µl, media + 1% PowerLoad + 0.1% Fluo-4 AM, #F10489, Thermofisher) was pipetted in chambers 1 and 3. Then chips, covered with aluminum foil, was placed in the incubator (37 0C) for 15 minutes. Sterile plastic adaptors, created by slicing along the upper marked sections of 1000 µL micropipette tips, were connected at inlets/outlets of the side channels (first and third channels only) **(Fig.2)**. Then, fluorescent microscopy (Leica DMI6000 Inverted, 10x objective) was used to record changes in calcium signaling intensities within the region of interest (ROI). Here the ROI was chosen to be a plane at ~100 µm from the bottom glass coverslip (approximate center plane of ~250 µm thick MLO-Y4-laden collagen in chamber 2). Before testing, the pump is connected to the chips’ inlet and outlets and left undisturbed for 15 minutes. For each calcium signaling experiment, data was captured at least 40 seconds before PUFFS was applied for 60 seconds at 0.33 Hz (or 1.66Hz), and imaging continued for another ~30 seconds after the end of PUFFS. Images were captured at the rate of 7 frames per second for total duration of ~2.5 minutes using LASX camera software. For experiments with two PUFFS applications, chips were subjected to PUFFS (0.33 Hz) followed by a rest period of 10-15 minutes before applying PUFFS at 1.66 Hz. This procedure was followed for both static and dynamic samples. For blocking experiments, 0.5 mg/mL of GAP26 (A1044, APExBIO, Cx43 gap junction blocker) was pipetted in chambers 1 and 3 for 45 minutes, washed 3x with media, before performing signaling experiments. All signal recordings were imported as LOF/LIF files into Image J. The measure function and the line tool were used to identify three sub-regions in chamber #2 based on the distance from PUFFS (Ch#3): proximal (0 - 285 µm), central (570 µm), and distal (855 µm). For all recorded timeframe-stacks, a z-projection mask with an outline of displacement of individual cells, and intensity within the masks were auto-traced using a specified threshold within the ROI manager. Using non-florescence regions (with no cells) as background, multi-measure option was used to analyze intensities within outlined regions for each timeframe slice, and data (mean, area, integrated density) was recorded in a CSV file spreadsheet. Oscillatory responses (responsive MLO-Y4s) were characterized under a specific threshold or number of peaks generated based on frequency conditions using the *PeakFinder*.*m* script run with MATLAB. The threshold peak ranged from 45 to 117 and 94 to 318 with 0.03 peak prominence, for frequencies 0.33 Hz and 1.66 Hz, respectively. Each response or column was sorted using the filter function, leaving oscillatory responses as the only response on the spreadsheet. The SORT function was also used to reorder responses based on the distance, grouping like cells together. Oscillatory cells within the same distance group were averaged into a single response; thus, each chip had 3 responses that correspond to each sub-region (proximal, central, distal). Individual intensities for each chip were plotted over time and *signalcharacteristics*.*m* script was used to compute the signaling frequency.

### RNA Harvest and RT-qPCR

PCR was performed with unstained replica sample batches. To assess the impact of PUFFS stimulation on gene expression by osteocytes, devices were subjected to Dynamic or Static conditions for 7 or 21 days. At the end of the study, the central channel containing the stimulated osteocytes was removed, snap frozen and stored at −80°C prior to RNA isolation. To isolate RNA, the isolated channels were homogenized in Trizol reagent (Thermo-Fisher, Grand Island NY) using a Precellys bead mill with MKR28 matrix (Bertin Technologies, Rockville MD) for three cycles of 30s at max speed. The RNA was extracted by the recommended protocol, and further purified using RNEasyPlus columns (Qiagen Inc., Valencia, CA). Integrity of extracted RNA was verified by Formaldehyde-Agarose electrophoresis (28S:18S rRNA >2:1), and RNA purity and quantity were assessed by UV spectrophotometry. The isolated RNA (35ng/sample) was then reverse transcribed to cDNA (Quantitect Reverse Transcription Kit, Qiagen) and amplified with (Quantitect SybrGreen PCR Kit, Qiagen) and oligonucleotide primers (**SI-6**; Azenta Life Sciences), on an Eppendorf Realplex2 instrument. A cDNA library prepared from mouse tibial bone tissue was used to verify primer specificity and optimize reaction efficiency. Dissociation curve analysis was used to verify reaction specificity for each reaction. Following qPCR, expression data normalized to the geometric mean expression of 2 housekeeping genes (*Gapdh* and *Hsp90ab*). Data are presented as mean fold difference ± 1SD (n=3-5 per group), relative to 7d static controls as calculated by −2^DDCT^ method. Statistical significance of differences between treatment groups was determined by 2-way ANOVA with time (7d vs 21d) and treatment (Static/PUFFS) taken as co-variates; Sidak post hoc test was used to evaluate pairwise differences between groups using GraphPad Prism Version 10.4.0 (527).

### Cell viability, morphology, connectivity plots, and immunofluorescence staining

Chips were stained with 0.05% calcein AM and 0.1% ethidium homodimer-1 and washed with PBS before imaging using Leica DMI6000 Inverted microscopy. For morphology assessment, chips were stained using 1 µg/mL DAPI (diamidino-2-phenylindole) for nucleus and 1:200 PBS-diluted Alexa Fluor™ Plus 488 Phalloidin (ThermoFisher) for f-actin, and imaged using upright Leica DM6 B fluorescent microscope equipped with THUNDER tissue imager and z-axis focal plane. To quantify cell number and connectivity within 3D MLO-Y4 networks, the following process was used. The 3D object counter in Image J was used to count the correct number of round cells for all time-points (time sequence images). The number of round cells obtained from the count mask was divided by the total number of cells (originally from the object mask). The 3D analysis tools gave output values for the x, y, and z locations for the cells using sliced files of the Nuclei-Blue channel. To determine the percentage of interconnectivity between timepoints, x and y values for each sample were imported onto a code-embedded Excel spreadsheet using the Gaussian-Euclidean distance equation. Once imported, distance measurements between cells were calculated with the code; here we assume that if the distance between two cells was ≤ 50 µm, the cells are connected to each other. For immunostaining, at specific time-points (Day 3, 7, and 21), chips were fixed (4% formaldehyde in PBS) for 15 minutes at room temperature, washed three times, followed by permeabilization using 0.2% for 10 minutes, and washing again for three times. Then, chips were incubated with 1% BSA (blocking agent) for 1 hour at room temperature, washed three times using PBS (15min for each wash). Chips were incubated with primary antibodies overnight at 4°C, then washed three times with PBS (15min for each wash) followed by incubation with secondary antibody solution. <u>Primary antibodies:</u> Mouse Monoclonal Connexin 43 antibody (#35-5000, Thermofisher) and Rabbit Polyclonal Integrin Alpha V + Beta 3 antibody (#BS-1310R, Thermofisher) or Sclerostin antibody (#219331AP, Thermofisher) at dilution of 1:200 in 0.2% BSA; 0.1% Tween; 0.3% Triton-X 100 (BTT). <u>Secondary antibodies:</u> Alexa Fluor Plus 647 goat anti-mouse IgG secondary antibody (#A32723TR, Thermofisher) and Alexa Fluor 594 Goat anti-Rabbit IgG (H+L) Highly Cross-Adsorbed secondary Antibody (#A32740, Thermofisher) at a dilution solution of 1:1000 in BTT. For control experiments, using Day 21 samples, the same procedure was followed in the absence of primary antibodies. The results show little-to-no non-specific fluorescence signals.

### Statistical Analysis

*O*ne-way and two-way ANOVA/Tukey tests were used to identify significant differences. For all results, ^*^ p ≤ 0.05; ^**^ p ≤ 0.010; ^***^ p ≤ 0.0010; ^****^p ≤ 0.0001 by 2-way ANOVA; Sidak post-hoc test used for pairwise comparisons.

## ACKNOWLEDGEMENTS

This work was financially supported by funding from the National Institutes of Health, R01 AR083466-01 to PS. We also acknowledge support from NIH COBRE Award 2P20 GM109024. Any opinions, findings, conclusions, or recommendations expressed in this material are those of the author(s)and do not necessarily reflect the views of the NIH or funding agencies

## Supplementary Materials

### SI-1

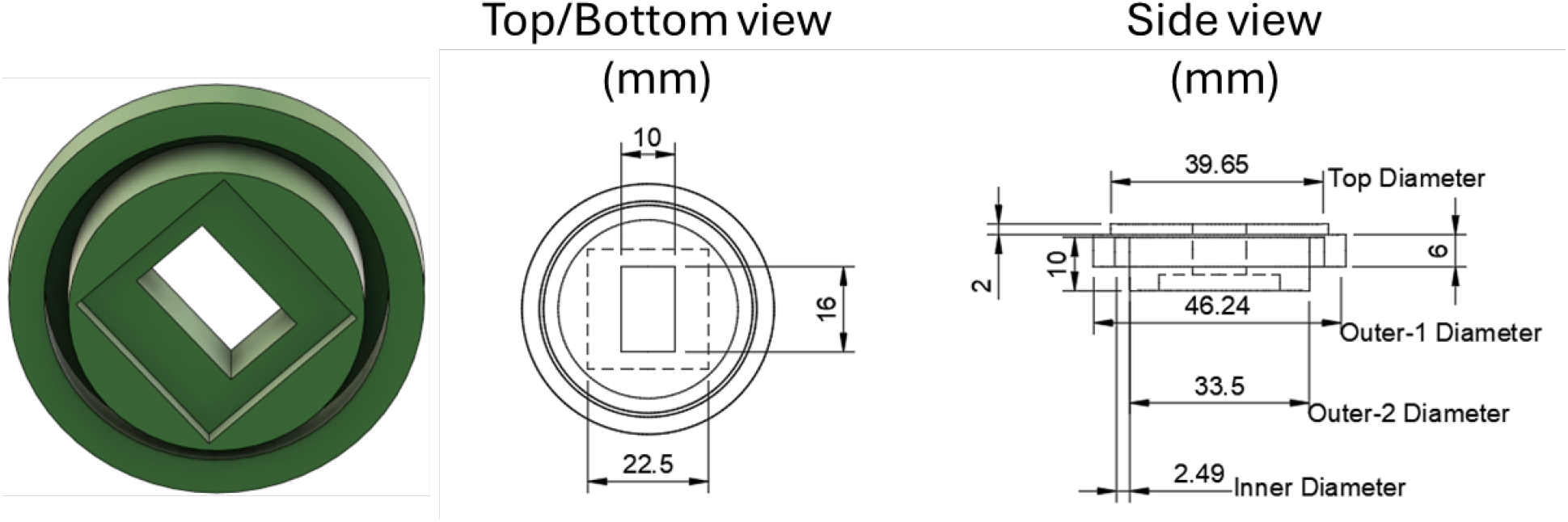
CAD schematic of device stabilizer with top, bottom, and side view dimensions in mm, generated by Fusion 360 drawing tool

### SI-2

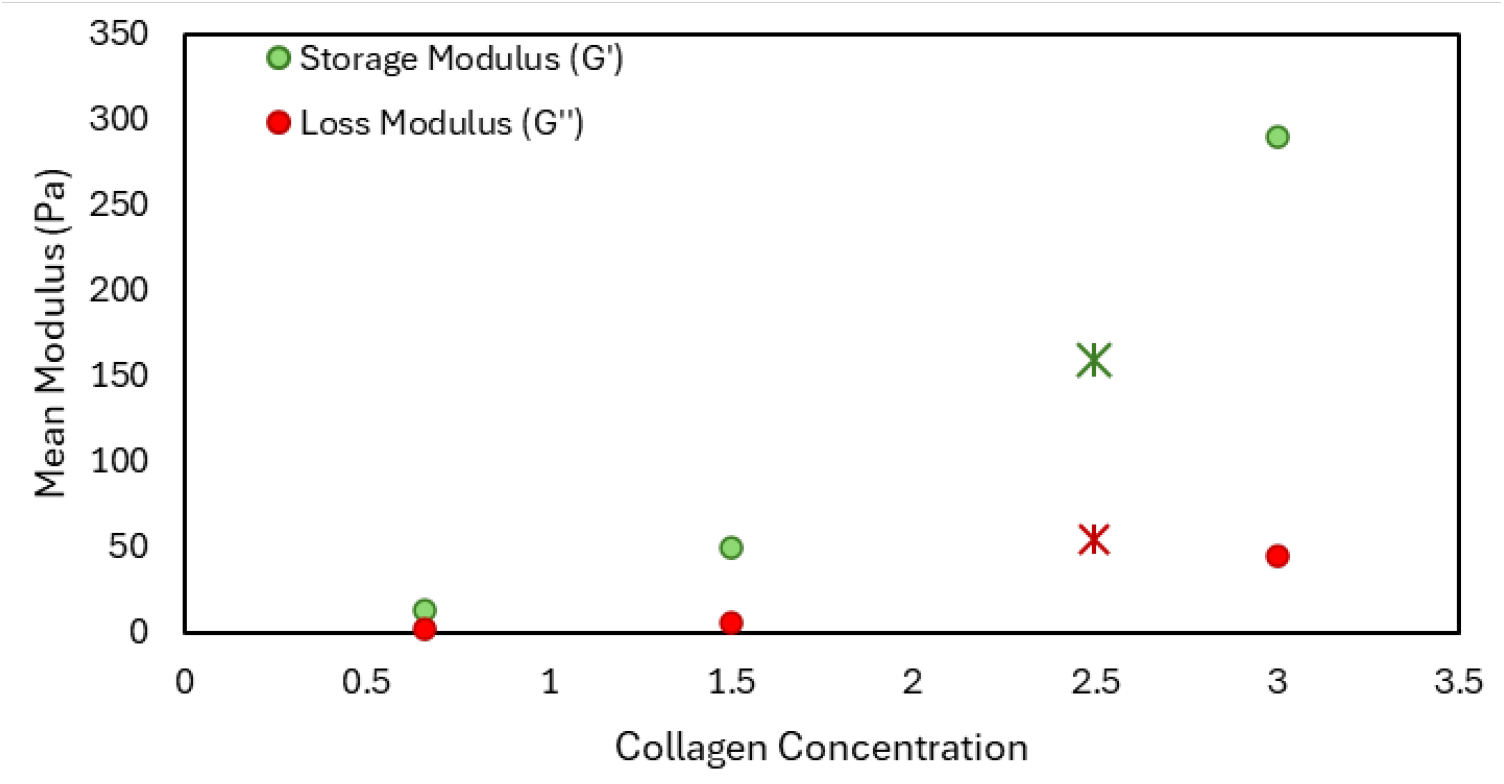
Mean modulus for Storage and Loss plotted for the experimental 2.5 mg/ml with literature values of 0.5, 1.5, and 3 mg/ml collagen. ***Reference:*** *Helary, Christophe, Isabelle Bataille, Aicha Abed, Corinne Illoul, Annie Anglo, Liliane Louedec, Didier Letourneur, Anne Meddahi-Pelle, and Marie Madeleine Giraud-Guille. “Concentrated collagen hydrogels as dermal substitutes*.*” Biomaterials 31, no. 3 (2010): 481-490*.

### SI-3

**Calculation of velocity and stress during application of PUFFS in chamber 3 of the PDMS microfluidic chip**.

Pulsating flow results in oscillating magnitudes of velocity and pressure. In a scenario where PUFFS forms a fluid layer bound by two parallel surfaces, we can determine an analytical solution to solve for

**Figure 1.**
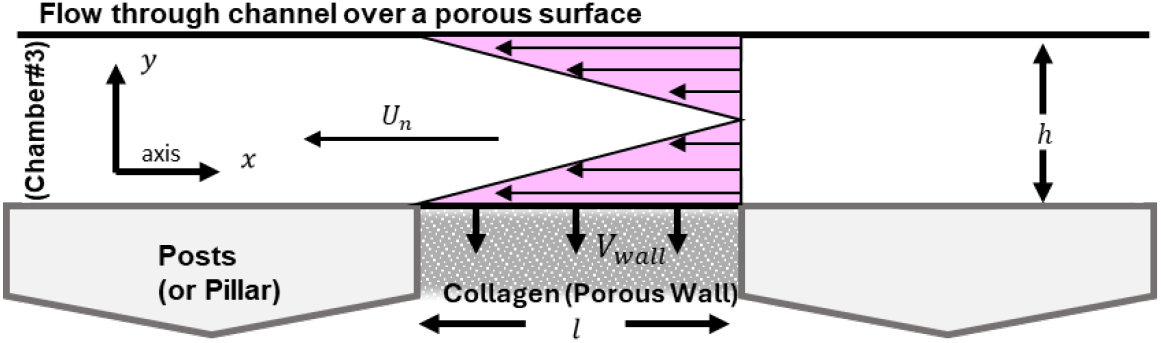
Cartoon schematic of shear stress along the collagen wall (or elastic region) at the gap regions of the channel where x-component velocity travels in the negative direction

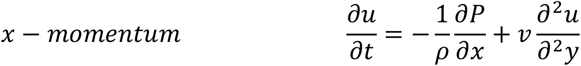

velocity that oscillates with time in the x direction. Assume fluid density is constant and no-slip conditions at the wall with boundary conditions *u*(*y* = 0) = 0 and *u*(*y* = *h*) = 0. The continuity equation simplifies to *dv*/*dy* = 0. The pressure gradient oscillates with time and can be derived as a Fourier series, 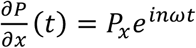 where ω is angular frequency for *n* harmonic. Further derivations will only consider the real part of the imaginary *i* complex. Thus, the steady-state solution can be represented as, *u*(*y, t*) = ∅(*y*)*e*^*inωt*^. We substitute the expressions into the x-momentum equation and rearrange them into second-order differential 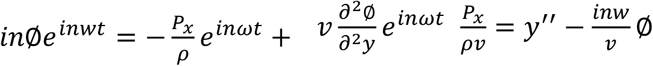. We satisfy the conditions of the homogenous equation on the left side to find the complementary solution, and then the particular solution to complete the general solution, 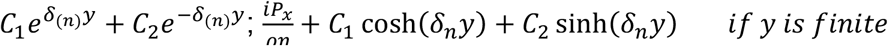 where, 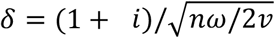 and *y* is finite. Applying the boundary conditions *u*(*y* = 0) = 0 which gives *C*_1_ = 0, and *u*(*y* = *h*) = 0 led to the solution for ∅(*y*) that gives the expression for *u*(*y, t*), the velocity of PUFFS through chamber #3; 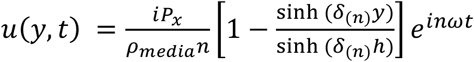

Collagen can be considered as a porous component. To determine shear stress at the wall, we may assume x-velocity is constant at these regions because the fluid covers the entire collagen surface giving no time for velocity or pressure variations. The continuity equation simplifies to *dv*/*dy* = 0 and the governing equation becomes 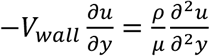 where, ∂*p*/∂*x* = 0.

By rearranging terms, we can integrate the second-order homogenous differential to find the following general solution: 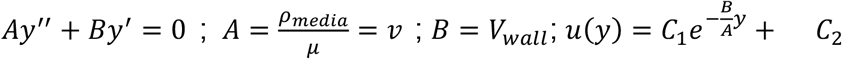. The boundary conditions *u*(*y* = 0) = 0 and *u*(*y* = *iinite*) = 0 give the exact solution for velocity in the fluid at the collagen region, followed by the shear stress at the wall by derivation 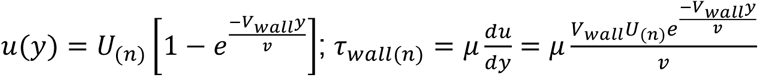; Here we can choose a *U*_(*n*)_ by plugging known parameters into the oscillating velocity profile (eq.6). The initial pressure (*P*_*n*=0_) in the x-direction was determined using Hagen-Poiseuille equation for rectangular channels, where *h* is width.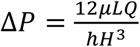; *P* = 36.80 *PP*; Pressure-driven flow through a porous medium can be calculated using Darcy’s Law. We assume that Darcy’s velocity is equivalent to the constant suction velocity into the collagen wall, which is near 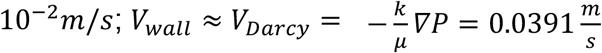. Deformation to the collagen wall is perpendicular to the length of the wall. The shear modulus ranged from 61 kPa to 42 kPa, using the following parameters from Table. Finally, to determine the maximum shear stress inflicted along the wall at the collagen surface, we used the computed values of *U*_*n* =1_ and 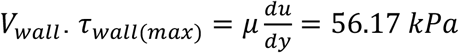

**Figure 1.**
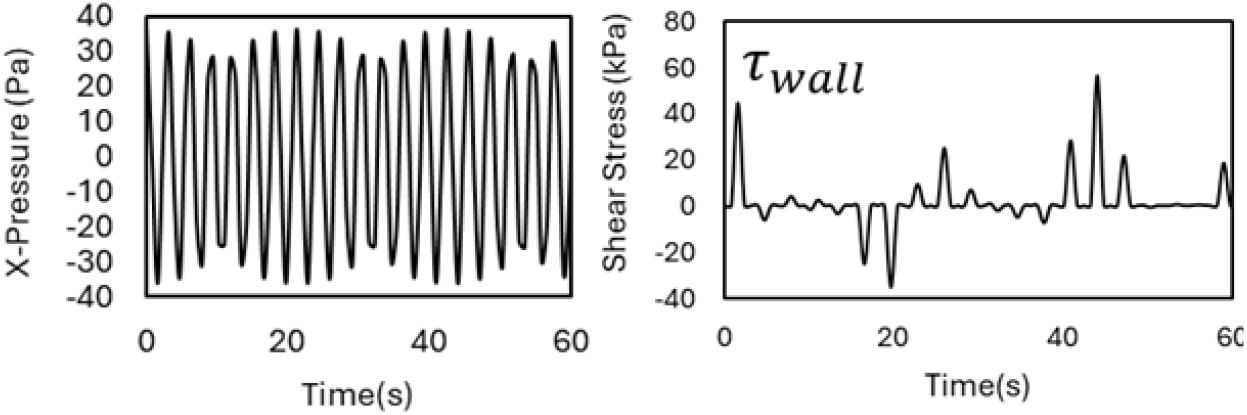
(Left) Plotted oscillating pressure and shear stress (right) along the wall at the collagen surface

### Relevant References

I.G. Currie. Pulsating Flow Between Parallel Surfaces. In: Faulkner LL, editor. Fundamental Mechanics of Fluids. Third, 2003; Recktenwald, Steffen M., Christian Wagner, and Thomas John. “Optimizing pressure-driven pulsatile flows in microfluidic devices.” Lab on a Chip 21, no. 13 (2021): 2605-2613.; Pérez-Rodríguez S, Huang SA, Borau C, García-Aznar JM, Polacheck WJ. Microfluidic model of monocyte extravasation reveals the role of hemodynamics and subendothelial matrix mechanics in regulating endothelial integrity. Biomicrofluidics 2021;15

**Table** Parameters used for analytical solution of velocity and pressure profiles in the x-direction

| Parameters | Notation | Value | Units |
| --- | --- | --- | --- |
| Length of collagen wall | $l$ | 6.50E-05 | m |
| Length of channel in ROI | $L$ | 2.80E-04 | m |
| Height in y | $h$ (width) | 5.13E-04 | m |
| Height in z | $H$ | 2.80E-04 | m |
| Cross-sectional area | $A$ | 1.44E-07 | m<sup>2</sup> |
| Viscosity media | $\mu$ | 8.89E-04 | Pa/s |
| Viscosity collagen | $\mu$ | 1.425 | kg/ms |
| Density (media) | $\rho_{media}$ | 1000 | kg/m<sup>3</sup> |
| Permeability (collagen)[4] | $k$ | 1E-07 | m/s |
| kinematic viscosity (media) | $\nu$ | 8.89E-07 | m<sup>2</sup>/s |
| Omega | $\omega$ | 6.28 | radian |
| Velocity flowrate (pump) | $V$ | 0.12 | m/s |
| Oscillating Fluid Velocity wall, | $U_{(n=1)}$ | 0.018 | m/s |
| Velocity flowrate at wall, | $V_{wall}$ | 0.039 | m/s |
| Volumetric flowrate (pump) | $Q$ | 1.67E-08 | m<sup>3</sup>/s |

#### SI-4

Stress in a collagen layer due to capillary traction on the surface

**Figure 1.**
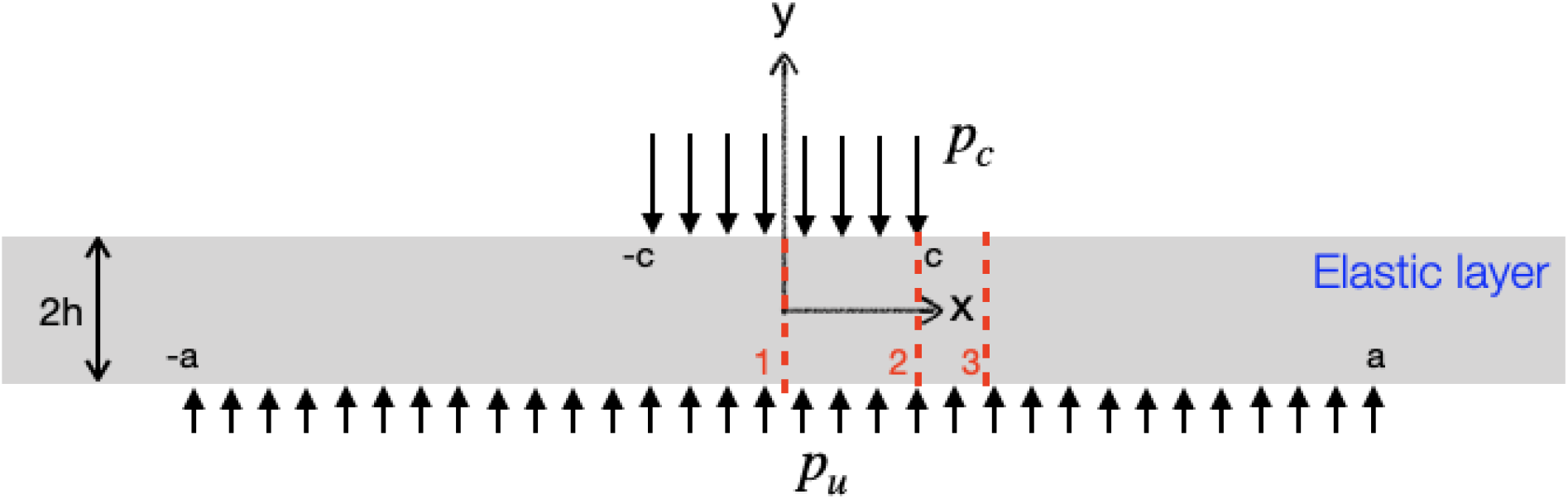
Geomtry of the problem studied here; an elastic layer of thickness 2*h* is subjected to a capillary pressure *p*_*c*_ = *γ/R* on the top over a width of 2*c*. A uniform pressure of *p*_*u*_ is acting on the bottom surface of the layer. Normal stress (*σ*_*yy*_) at the three sections (marked by the three red dashed lines) are shown in fig. 3.

Here we calculate the stress in a collagen layer which is subjected to capillary traction due to a steadily propagating droplet. We assume the collagen layer to be purely elastic, and the geometry is sketched in fig. 1. We consider the problem to be plain strain problem and the layer thickness (2*h*) to be much smaller than the lateral dimensions. On the top surface a capillary pressure is exerted from the moving droplet. We focus on the static problem with the capillary pressure on top to be stationary. The bottom surface of the elastic layer is exposed to an uniform pressure. Thus equilibrium requires that,

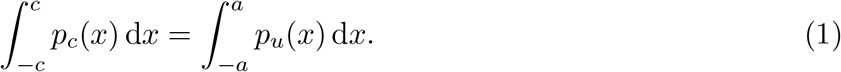

Since, *p*_*c*_ = *γ/R*, the above condition gives that *p*_*u*_ = *p*_*c*_(*c/a*) = *γ/R*(*c/a*). The equilibrium of of the elastic layer is given by,

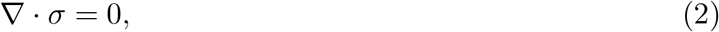

where *σ* is the stress tensor which has three independent components of *σ*_*xx*_, *σ*_*yy*_, and *σ*_*xy*_. Equation (2) is solved along with the boundary conditions, *σ*_*yy*_(*y* = *h*) = *−p*_*c*_, *σ*_*yy*_(*y* = *−h*) = *−p*_*u*_, *σ*_*xy*_(*y* = *±h*) = 0. We follow a Airy’s stress function formulation to solve the above equation. In this method, a scalar stress function, Φ is chosen such that, *σ*_*xx*_ = *∂*^2^Φ*/∂y*^2^, *σ*_*yy*_ = *∂*^2^Φ*/∂x*^2^, and *σ*_*xy*_ = *−∂*^2^Φ*/∂x∂y*. This choice ensures that Φ identically satisfies eq. (2). However, we are left with the unknown, Φ which is found from the compatibility equation which is the biharmonic equation

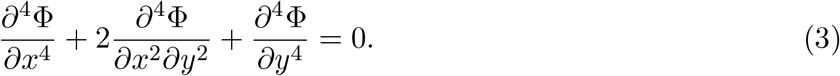

We solve the above equation using Fourier transform, which essentially transform the above PDE to an ODE, 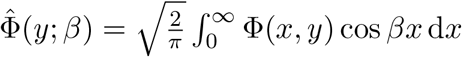,

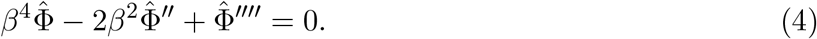

Here *β* acts as a parameter, and derivatives are takes with respect to *y*. It is to be noted that we have already taken into account the symmetry of the problem about *y*-axis in defining the Fourier transform using a cosine kernel. Eq. (4) has a general solution

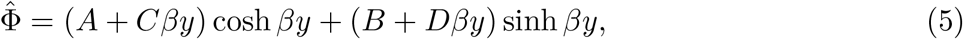

where *A, B, C*, and *D* are constants which are found from the boundary conditions. Thus we transform the boundary conditions above in the Fourier domain to find,

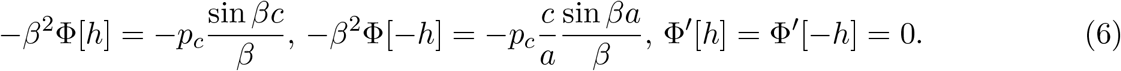

Using these boundary conditions, we find the four constants as

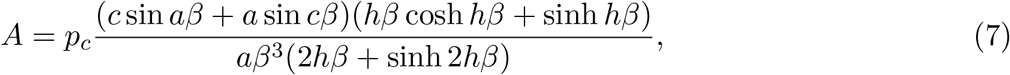

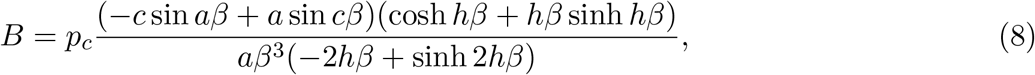

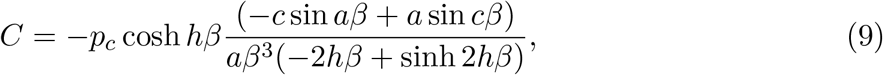

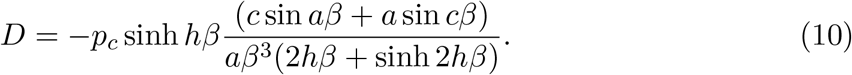

Thus we find an analytical solution of the Airy’s stress function in Fourier domain. However, the stress function and subsequently the stress components in terms of x and y coordinates are found by numerically evaluating the inverse transform,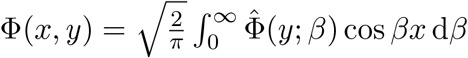.

Since the forces acting on the elastic layer is purely compressive, the dominant stress component is the normal stress along y, *σ*_*yy*_. In the following, we show how *σ*_*yy*_ varies across the elastic layer. For this purpose, we scale all the lengths by *h*, and stress by *p*_*c*_. Figure 2 shows the distribution of *σ*_*yy*_*/p*_*c*_ in the elastic layer around the droplet. This plot is obtained for *c/h* = 2 and *a/h* = 10, and *p*_*u*_*/p*_*c*_ = *c/a* = 1*/*5. Note that, the stress is maximum at the top surface, beneath the droplet reaching a value of 1 which is marked by color blue. While outside the droplet, the top surface is stress free as marked by the red color. The orange represents the compressive stress at the bottom surface.

**Figure 2.**
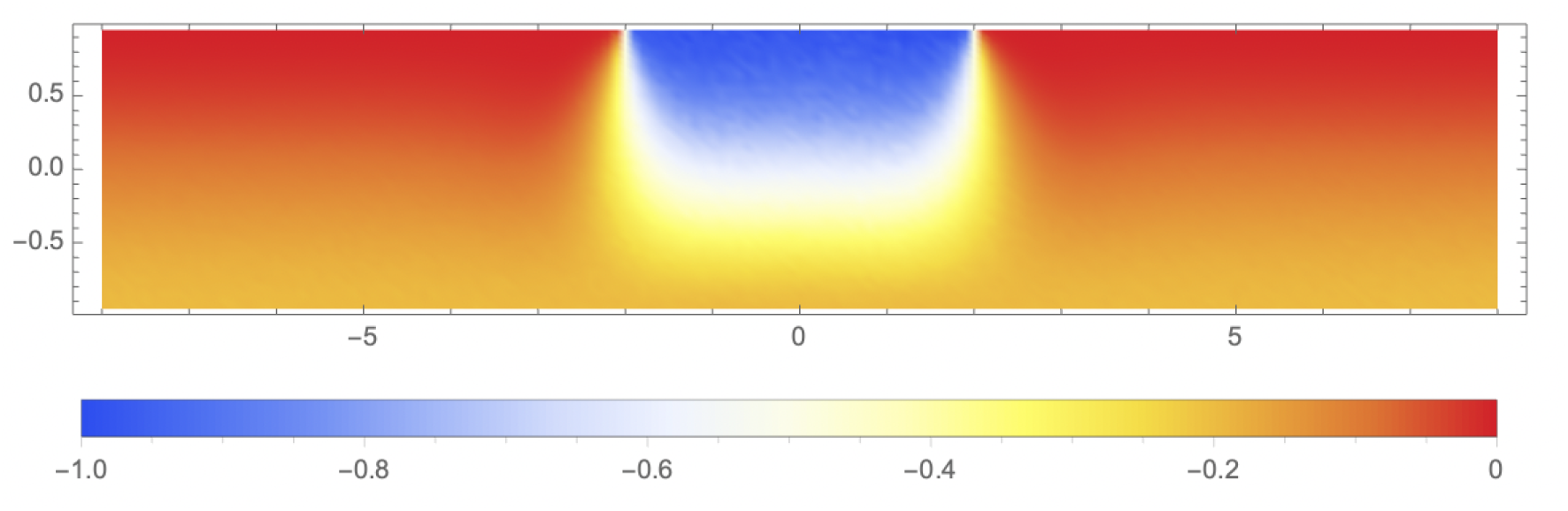
Colormap showing distribution of *σ*_*yy*_*/p*_*c*_ in the collagen layer.

Now we plot the variation of *σ*_*yy*_*/p*_*c*_ with *y/h*. For this, we consider three cross-sections marked by the red, dashed lines in fig. 1. Figure 3 shows how this dimensionless stress varies along the depth of the elastic layer at the three sections

**Figure 3.**
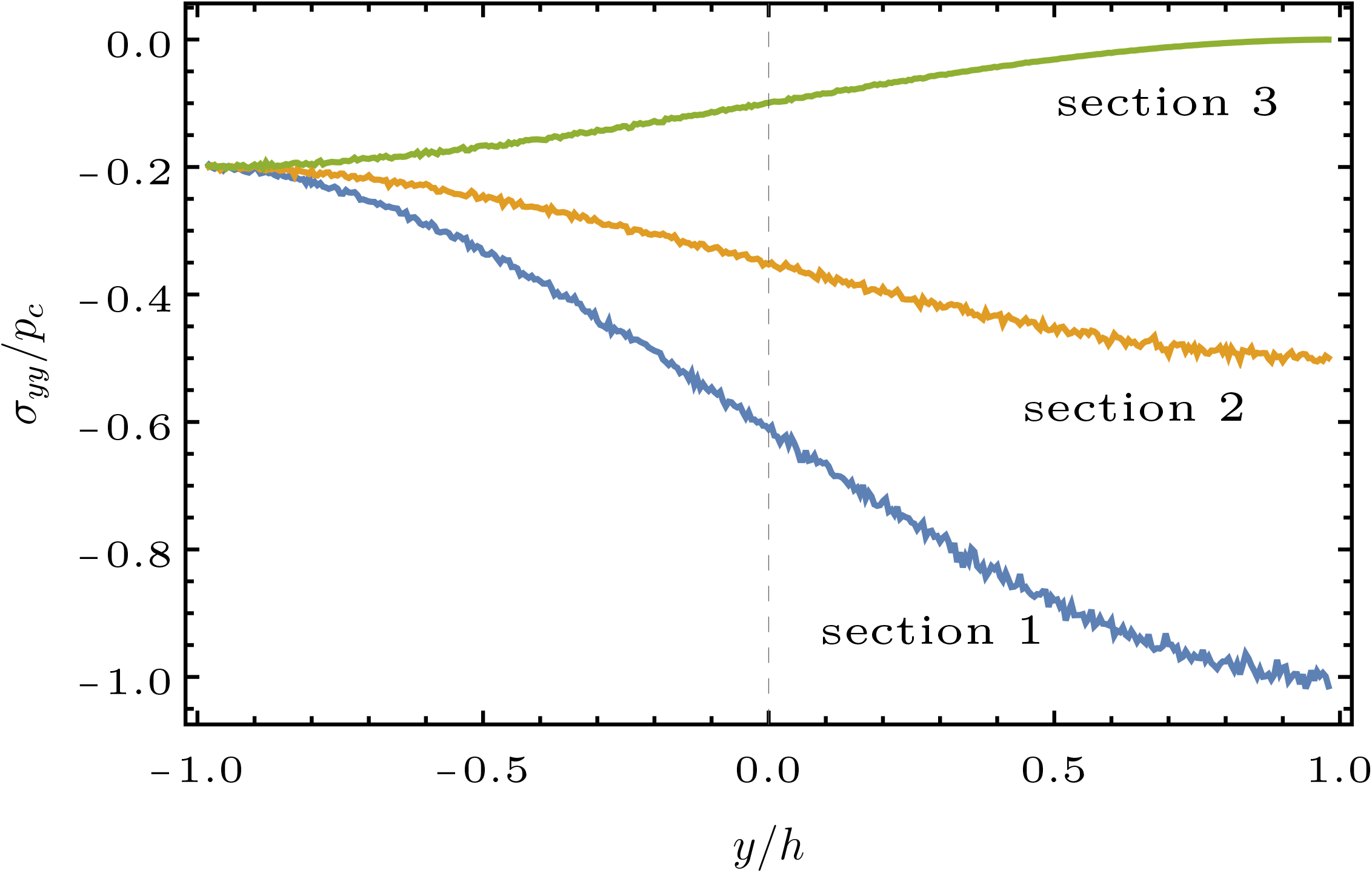
Variation of *σ*_*yy*_*/p*_*c*_ across the thickness of the elastic layer.

#### SI-5

To study the dynamic interactions between water droplets and a collagen medium, we developed a Eulerian viscous two-phase model to simulate the velocity distribution within the collagen gel, treating water droplets as the primary phase and the collagen gel as the secondary phase. The water droplets were assumed to undergo a combination of rotational and translational motion, and surface tension forces were considered to capture the interactions at the interface between the water and collagen gel phases. The simulation results revealed high-velocity regions near the interface between the water droplets and the collagen gel, with a subsequent decline in velocity within the bulk due to the gel’s viscous resistance. By comparing the experimental velocities at specific points within the collagen medium to the simulated values, we identified correction factors that aligned the numerical predictions with the experimental data, thereby validating our simulation model.

### Numerically modeled velocity profile in the collagen medium

Through the in silico architecture shown in figure 1a, we have run a two-phase flow simulation to numerically model the velocity field response in the collagen medium (see figure 2b) owing to the interfacial interactions with the water drops moving through the side channel (labeled as chamber 1 in figure 1a). The velocity contour plot in the collagen bulk (marked as chamber 2 in figure 1a) reveals a high-velocity region near the interface between the drop and the collagen gel, at the sharp channels that cap the grooved enclosure for chamber 2. Moving cross-stream (relative to chamber 1), note that the negative sign on the velocity values implying the collagen bulk velocity is directed along the negative *y*-axis. The maximum velocity at the inlets is approximately −7.03 × 10^−2^ m/s, as shown in the color scale. The water interacts with the collagen gel, creating this higher velocity region near the interface. Subsequently, in the yellow and green regions close to the inlets, the velocities range approximately from −2.30 × 10^−2^ to −4.66 × 10^−2^ m/s. The velocity magnitude declines progressively as we move further into the bulk owing to the collagen gel’s viscous resistance, with the least velocity approximating −0.53 × 10^−2^ m/s. Given the “barriers” in the contour plots, the velocity distribution results can be mathematized using a piece-wise function to describe the *y*-velocity profile of the collagen gel. We define the velocity *V*_*y*_ as a function of *y* along the vertical direction. From the observations of the contour plot, we can divide the domain into three major regions, namely:

- Inlet region: High-velocity region at the entry locations.
- Interface region: Region where water interacts with the collagen gel.
- Deceleration region: Region where velocity decreases owing to viscous resistance.

Assuming *y*_1_ as the boundary between the inlet and interface regions, *y*_2_ as the boundary between the interface and deceleration regions.

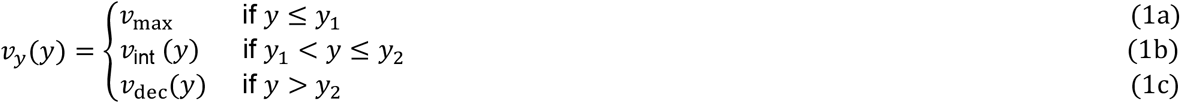

where *v*_ma*x*_ ≈ −7.03 × 10^−2^ m/s at the inlets, *v*_int_ (*y*) is the velocity in the interface region, which decreases from *v*_ma*x*_ to *v*_dec, max_ and *v*_dec_(*y*) is the velocity in the deceleration region, which further decreases to the minimum simulated velocity. For simplicity, assuming a linear decrease in velocity in the interface region.

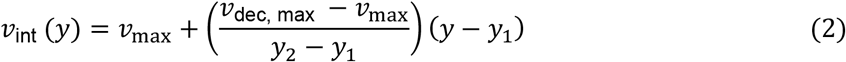

Where *v*_dec,ma*x*_ ≈ −2.30 × 10^−2^ m/s. Again, considering a bulk linear decrease in the deceleration region.

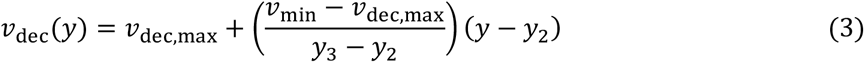

where *v*_min_ ≈ −0.53 × 10^−2^ m/s is the minimum velocity observed, and *y*_3_ is the vertical extent of the domain. Combining these equations, we get the piecewise function for *v*_*y*_(*y*) as:

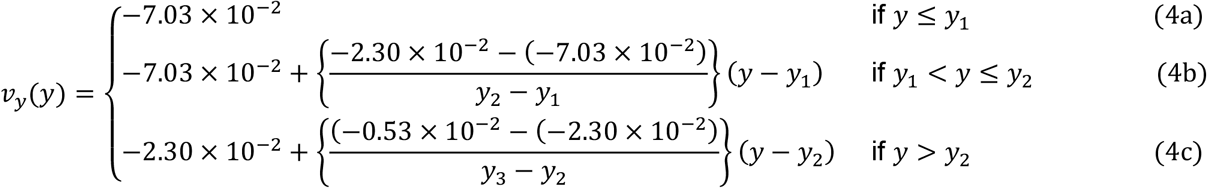

The piece-wise functions mathematically summarize the velocity distribution processed in the simulation results.

### Correction factor analysis for velocity discrepancies

The above model findings assume the water drops to be undergoing pure rotation through chamber 1 (see methods) and consequently over-estimate the momentum transferred to the collagen gel. In reality, the drop motion is a mix of rotation and translation along with viscoelastic stretching in presence of micro-scale surfactants, and as such a correction factor is essential to compare and analyze the velocity data from the numerical simulations and the experimental observations. The experimental velocities at two specific points (see figure 1b), the first point (#1) and second point (#2), were measured to be 9.91 × 10^−5^ m/s and 6.17 × 10^−5^ m/s. These velocities were determined by tracking the vertical displacement over 27 frames, providing an overall measure of the bulk movement at those points. The total displacement was divided by 2 to account for the oscillatory movement in both the positive and negative *y*-directions, which is assumed to be symmetrical owing to the periodic flow against the collagen wall. Correspondingly, the simulated absolute velocity regions for the two sample points were 1.12 × 10^−2^ m/s and 0.53 × 10^−2^ m/s respectively. The ratio of the velocities, projected numerically and observed experimentally, are hence of the same order, 𝒪(0). Be as it may, given the limitation of the model assumptions, we invoke a correction factor to account for the discrepancies in velocity measurements in the collagen medium. By calculating the ratio of experimental to simulated velocities for each point, we obtain the correction factors, respectively for #1 and #2 as:

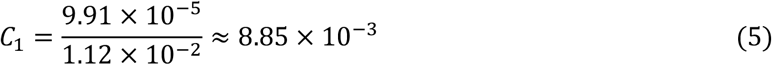

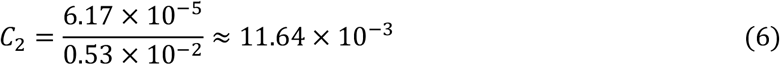

The correction factors are hence of the same order, justifying the relevance of the numerical findings when the simulated flow field is revised with a consistent correction coefficient applied across the domain. More importantly, the spatial demarcations of the simulated flow barriers in the collagen medium match almost exactly with the experimental trends.

To further refine our study and account for the complex interactions within the collagen medium, we introduce an elasticity model. This model provides a physics-based rationale for the correction by incorporating the elastic resistance from the gel layers, which the viscous model alone cannot fully capture. Specifically, the elasticity model examines the distribution of stress and the resultant deformation within the collagen layer when subjected to capillary traction forces. By analyzing the stress response in the collagen layer due to capillary forces from a steadily propagating droplet, we can better comprehend the dynamic interactions.

Here we calculate the stress in a collagen layer which is subjected to capillary traction due to a steadily propagating droplet. We assume the collagen layer to be purely elastic, and the geometry is sketched in fig. 3. We consider the problem to be plain strain problem and the layer thickness (2*h*) to be much smaller than the lateral dimensions. On the top surface a capillary pressure is exerted from the moving droplet. We focus on the static problem with the capillary pressure on top to be stationary. The bottom surface of the elastic layer is exposed to an uniform pressure. Thus equilibrium requires that,

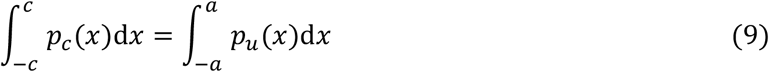

Since, *p*_*c*_ = *γ*/*R*, the above condition gives that *p*_*u*_ = *p*_*c*_(*c*/*a*) = *γ*/*R*(*c*/*a*). The equilibrium of of the elastic layer is given by,

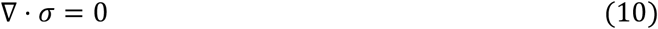

where *σ* is the stress tensor which has three independent components of *σ*_*xx*_, *σ*_*yy*_, and *σ*_*xy*_. Equation (8) is solved along with the boundary conditions, *σ*_*yy*_(*y* = *h*) = −*p*_*c*_, *σ*_*yy*_ (*y* = −*h*) = −*p*_*u*_, *σ*_*xy*_(*y* = ±*h*) = 0. We follow a Airy’s stress function formulation to solve the above equation. In this method, a scalar stress function, Φ is chosen such that, *σ*_*xx*_ = ∂^2^Φ/ ∂*y*^2^, *σ*_*yy*_ = ∂^2^Φ/ ∂*x*^2^, and *σ*_*xy*_ = − ∂^2^Φ/ ∂*x* ∂*y*. This choice ensures that Φ identically satisfies eq. (8). However, we are left with the unknown, Φ which is found from the compatibility equation which is the biharmonic equation

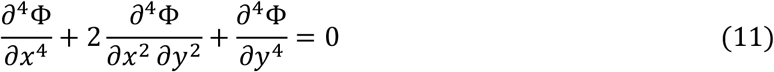

We solve the above equation using Fourier transform, which essentially transform the above PDE to an ODE, 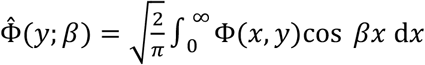,

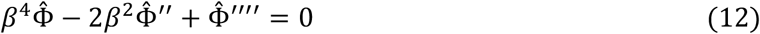

Here β acts as a parameter, and derivatives are takes with respect to *y*. It is to be noted that we have already taken into account the symmetry of the problem about *y*-axis in defining the Fourier transform using a cosine kernel. Eq. (10) has a general solution

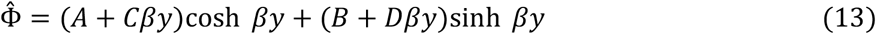

where *A, B, C*, and *D* are constants which are found from the boundary conditions. Thus we transform the boundary conditions above in the Fourier domain to find,

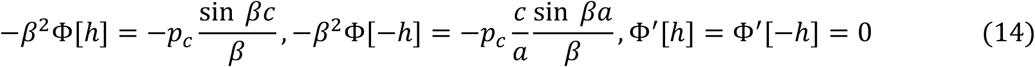

Using these boundary conditions, we find the four constants as

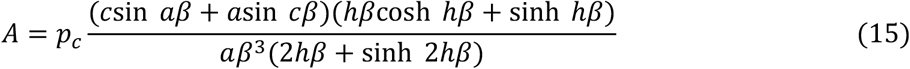

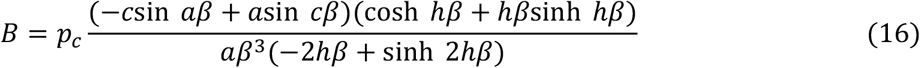

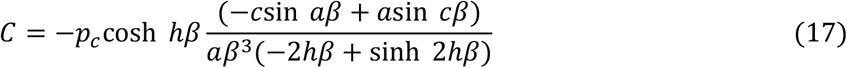

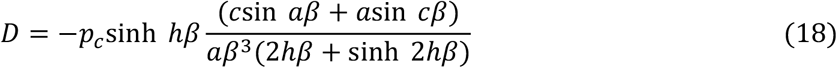

Thus we find an analytical solution of the Airy’s stress function in Fourier domain. However, the stress function and subsequently the stress components in terms of x and y coordinates are found by numerically evaluating the inverse transform,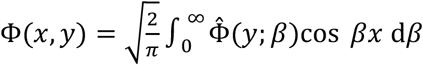.

Since the forces acting on the elastic layer is purely compressive, the dominant stress component is the normal stress along y, *σ*_*yy*_. In the following, we show how *σ*_*yy*_ varies across the elastic layer. For this purpose, we scale all the lengths by *h*, and stress by *p*_*c*_. Figure 4 shows the distribution of *σ*_*yy*_/*p*_*c*_ in the elastic layer around the droplet. This plot is obtained for *c*/*h* = 2 and *a*/*h* = 10, and *p*_*u*_/*p*_*c*_ = *c*/*a* = 1/5. Note that, the stress is maximum at the top surface, beneath the droplet reaching a value of 1 which is marked by color blue. While outside the droplet, the top surface is stress free as marked by the red color. The orange represents the compressive stress at the bottom surface.

### On the limitations in the numerical model for inter-phase interactions

The numerical simulations offer valuable insights into the interactions between water and collagen gel, but several limitations exist. The simulations used a planar representation of the experimental design, which does not fully capture the three-dimensional nature of the actual system. Additionally, the material properties used in the simulation, such as the viscosity and density of the collagen gel, were based on literature reviews and may not accurately reflect realistic variations due to factors like gel concentration and temperature. In the experiment, both air and water were present at the inlets. However, the simulation only considered water, as air was not the primary concern in studying the interaction between water and collagen gel. Moreover, the simulation did not account for the full rotational motion of water drops, requiring a correction factor to align the simulated velocities with the experimental measurements (see results).

## Materials and Methods

### In silico test geometry and spatial discretization

The two-dimensional structure of the experimental setup is illustrated in figure 1a, comprising three chambers, with chambers 1 and 3 being identical, bearing heights *h* = 0.5131 mm, while chamber 2 has a height *H* = 0.7980 mm. In figure 1a, the streamwise length *l* = 2.065 mm. Water drops with diameters *D* = *h* are modeled to be transiting through chamber 1. The geometry in panel (b) was spatially meshed, resulting in 47,740 linear quadratic elements. This design includes seven inlets for water entry, with the entire chamber 2 filled with collagen gel.

### Numerical simulation of interfacial mechanics-induced bulk motion in the collagen medium

The interaction of water (phase 1) with the surface of collagen gel (phase 2) through the inlets is modeled as a viscous laminar transient flow with SIMPLEC pressure-velocity coupling and second-order upwind spatial discretization. To replicate this interaction, the Eulerian multiphase model is used between the primary and secondary phases ^1^. This Eulerian multiphase model tracks the continuity and momentum for each phase ^2^. The simulation begins with the continuity equation ^3^, which ensures the mass conservation of each phase, mathematically implying

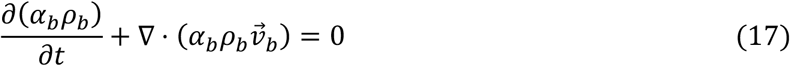

Here 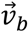 represents the velocity of phase *b, α*_*b*_ denotes the volume fraction, and *ρ*_*b*_ is the density of phase *b*. By adhering to equation 7, the simulation ensures that the mass of the water and collagen gel is conserved within the computational domain. This conservation also means that there are no sources or sinks in the system. The surface tension force ^4^ is considered to model the interaction at the interface between the water and collagen gel phases, and is quantified as

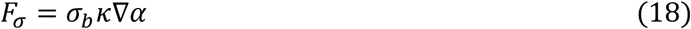

where *σ*_*b*_ is the surface tension coefficient, *κ* is the curvature of the interface, and ∇*α* is the gradient of the volume fraction. The curvature *κ*^3^ of the interface is calculated as:

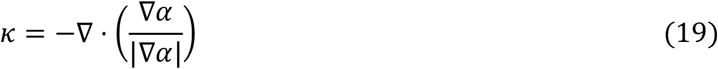

The above equations capture the dynamic behavior of the interface between the water and collagen gel phases. Subsequently, the momentum equation ^5^ describes the forces acting on each phase. The conservation of momentum principle, which is fundamental for determining the velocity and pressure fields within the system, is mathematized as

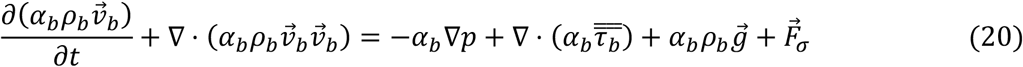

In this equation, *p* represents the pressure shared by both phases, 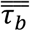 is the stress-strain tensor, and 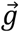 denotes the gravitational acceleration. This comprehensive equation incorporates the effects of pressure gradients, viscous stresses, gravitational forces, and surface tension, providing a detailed description of the flow dynamics within the system. The stress tensor for phase *b* is formulated to capture the viscous behavior and is given by ^6^

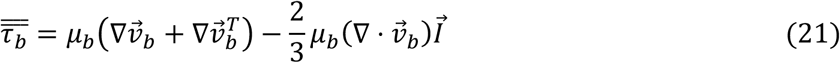

with *µ*_*b*_ being the dynamic viscosity, 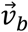 as the velocity vector, 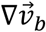 as the velocity gradient tensor, 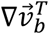 as the transpose of the velocity gradient tensor, 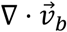 as the divergence of the velocity field, and 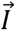 as the identity matrix. The above equation represents the stress distribution within the system. Finally, the volume fraction ^7^ for each phase is tracked using the volume fraction equation which monitors how the water phase infiltrates and interacts with the collagen gel, and is given by

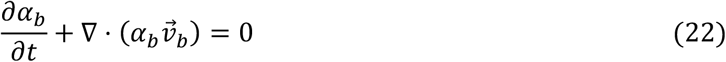

The convergence of the numerical solution is determined by minimizing the residuals of the mass and velocity components. For the simulations of pressure-gradient-driven laminar flow, typical execution times range from 0.2 to 0.5 hours for 500 iterations with a time-step of 0.001 seconds, utilizing a 4-processor-based parallel computation setup operating at 3.1 GHz on Xeon nodes. Assuming pure rotational motion of the water drops through chamber 1, the inlet velocity (at the water-collagen interface) was periodically specified by a user-defined function, bearing the following magnitude in the negative *y*-direction (see figure 1):

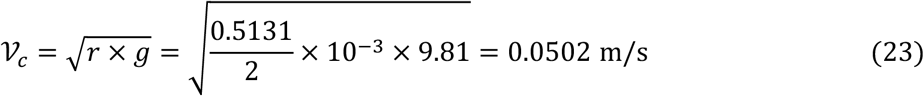

where *r* is the radius of the water drop and *g* is the gravitational acceleration. Physical parameters such as the density and viscosity of water are assumed to be 1000 kg/m^3^ and 0.001 kg/(m. s), respectively. For collagen gel, the density is assumed to be 1300 kg/m^3 8^ and viscosity 1.425 kg/(m. s)^9^.

### Figures and Tables

**Figure 1.**
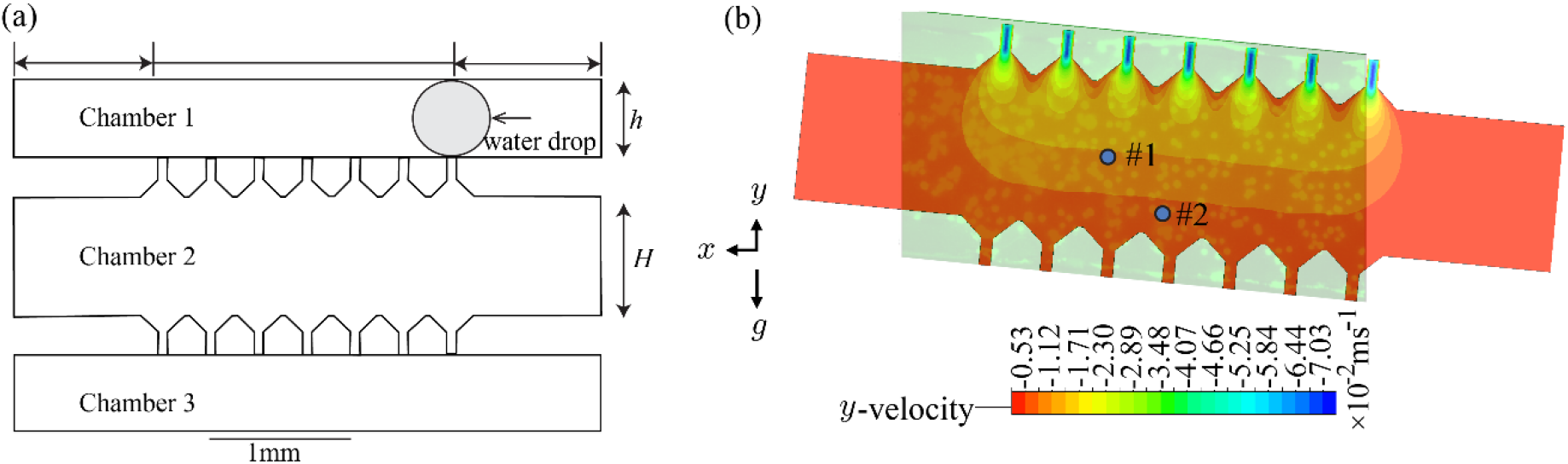
1 Panel (a) shows the planar numerical domain, extracted as a projection of the experimental setup. Panel (b) compares the experimental and numerical results through superposition, showing two points (#1) and (#2) in the numerical velocity profile used to calculate the correction factor (see results) where the color scale presents the *y*-component of velocity in units of 10^−2^ m/s, with dark blue indicating higher downward velocities and red indicating lower downward velocities. The dark red sections at the edges indicate places where the vertical velocity approaches zero (no-slip). The negative sign before the velocity indicates the velocity directed towards negative y-axis. The 1-mm scale reference bar is at the bottom of panel (a). Additionally, next to panel (b), *x* and *y* axes establish the spatial orientation and *g* signifies the gravity direction in the simulation.

**Figure 2.**
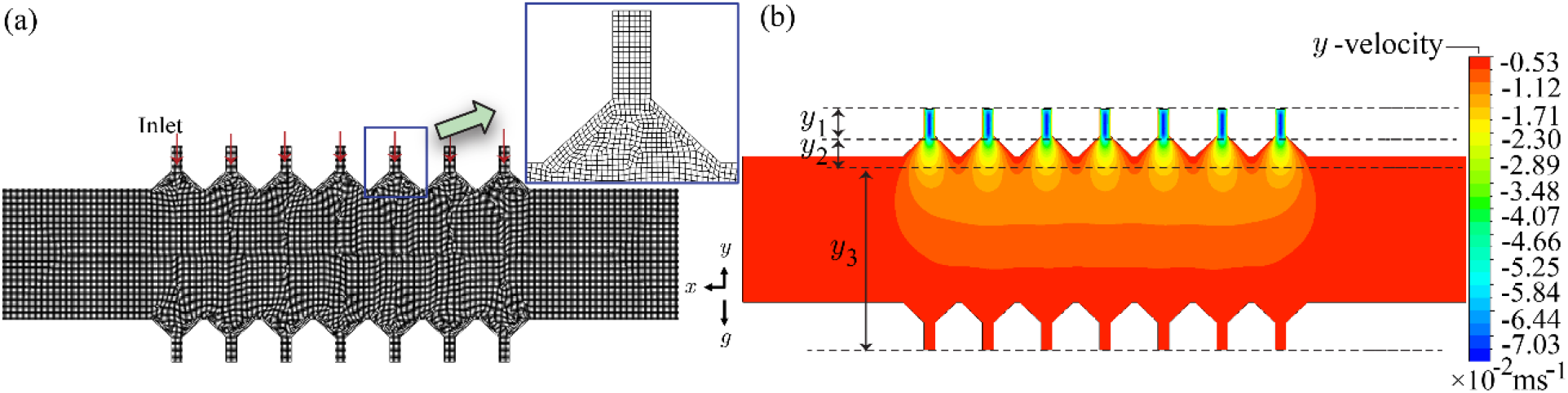
Panel (a) shows the mesh visual and its zoomed-in details, with the enforced inlets marked. Therein the red arrows show the direction of inlet flow. Panel (b) presents the velocity magnitude contours in the collagen gel, where the color scale presents the *y*-component of velocity in units of 10^−2^ m/s, with dark blue indicating higher downward velocities and red indicating lower downward velocities. The dark red sections at the edges indicate places where the vertical velocity approaches zero (no-slip). The negative sign before the velocity indicates the velocity directed towards negative y-axis. Additionally, next to panel (b), *x* and *y* axes establish the spatial orientation and *g* signifies the gravity direction in the simulation.

## SI-6

**Table. Oligonucleotide Primers.**

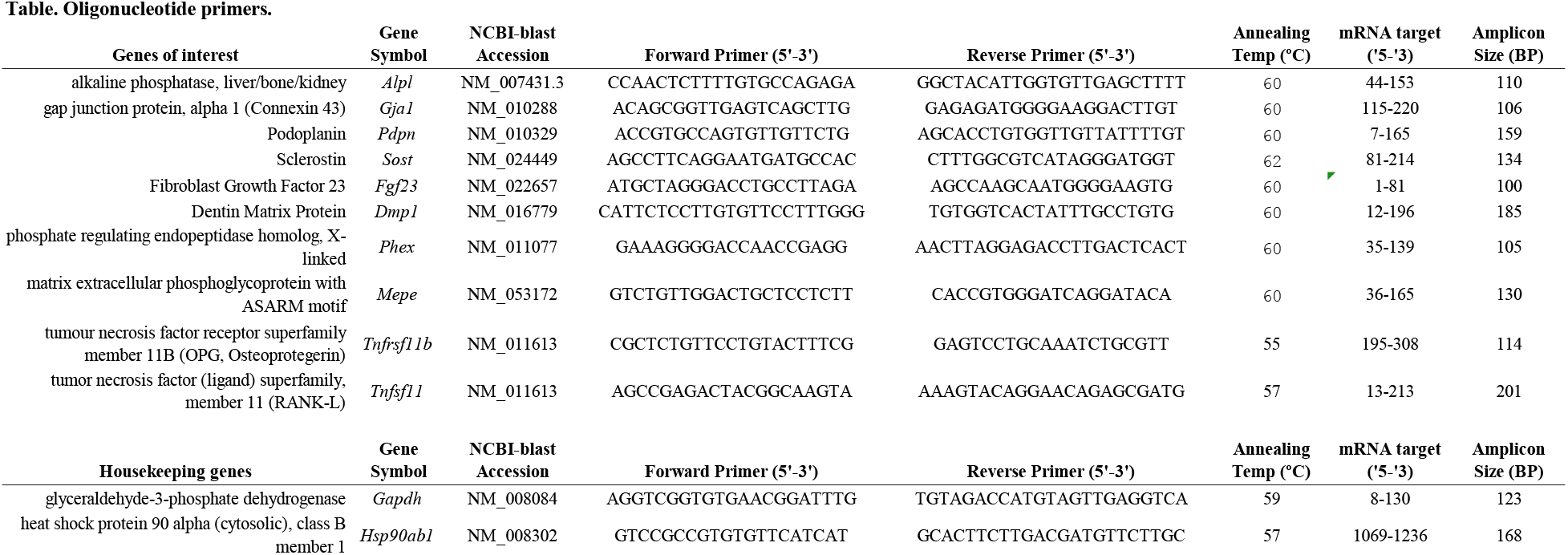

**Table Relative fold-change of gene expression by RT-qPCR**

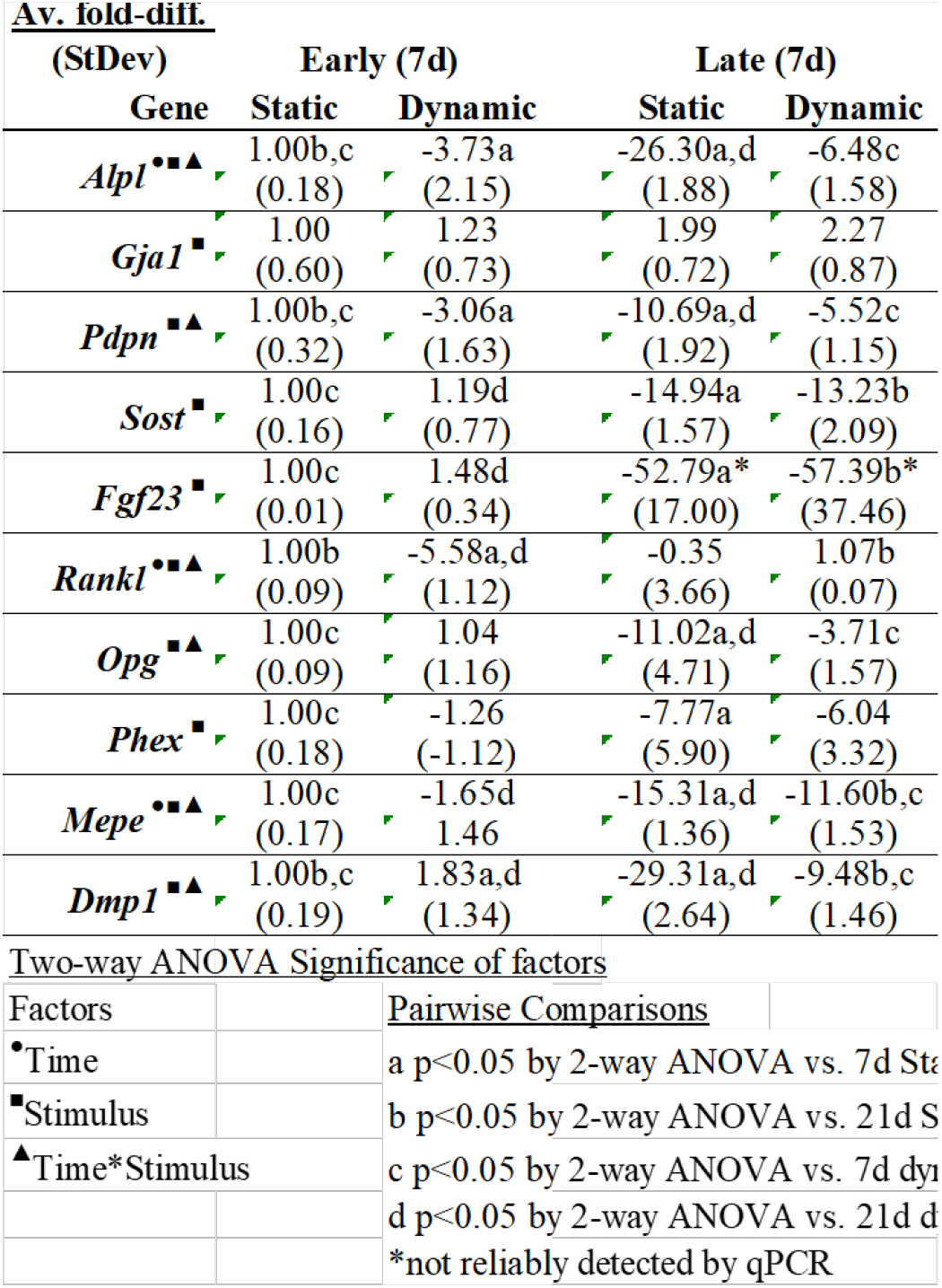

## SI-7

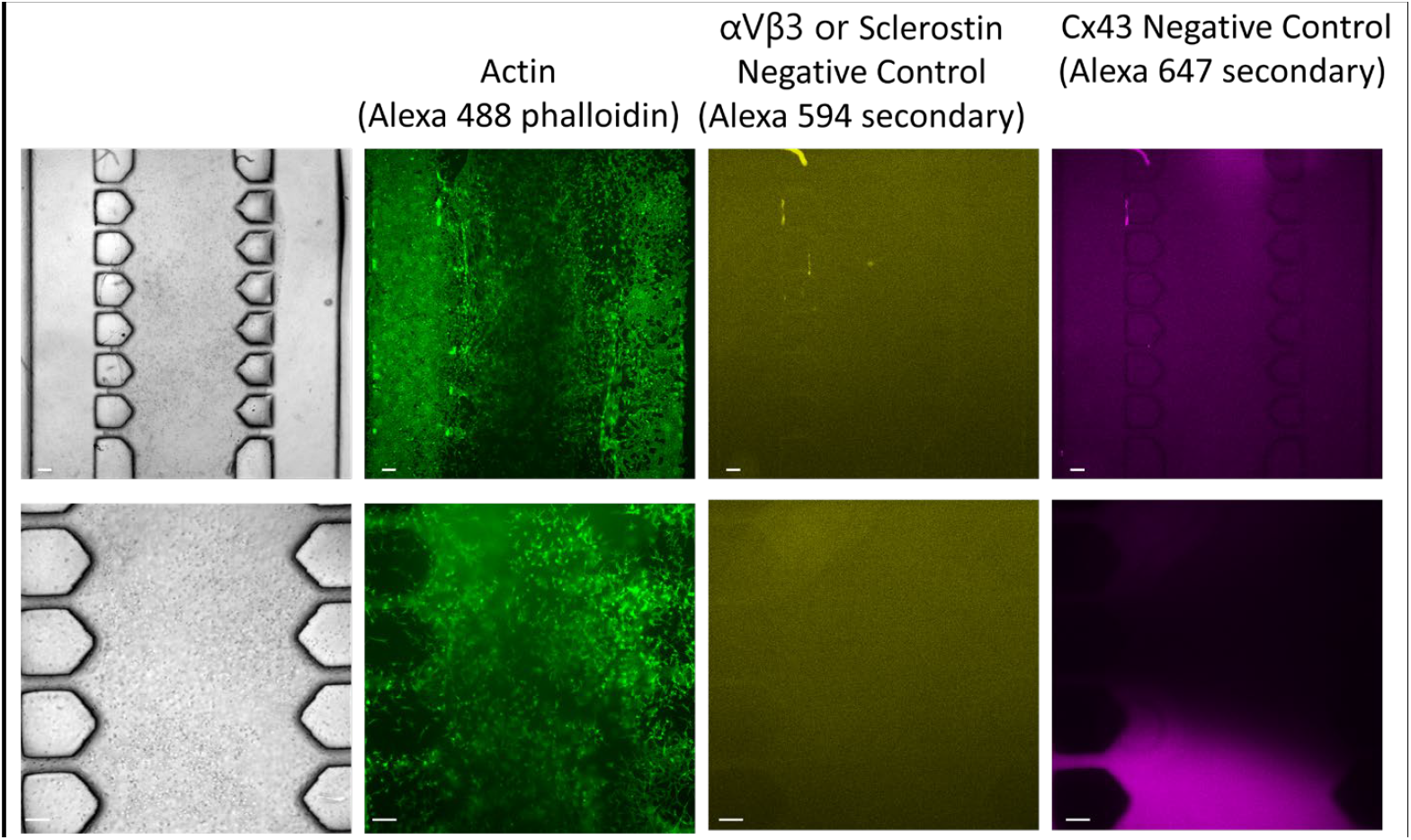
Control experiment of secondary antibody stains to show non-specific binding using Day 21 samples. Brightfield image of the sample to the far left. Actin was imaged under the green channel, Alexa 594 secondary was imaged under the red channel (depicted as a yellow hue signal), and Alexa 647 secondary was imaged under the far-red channel (depicted as a purple

## SI-8

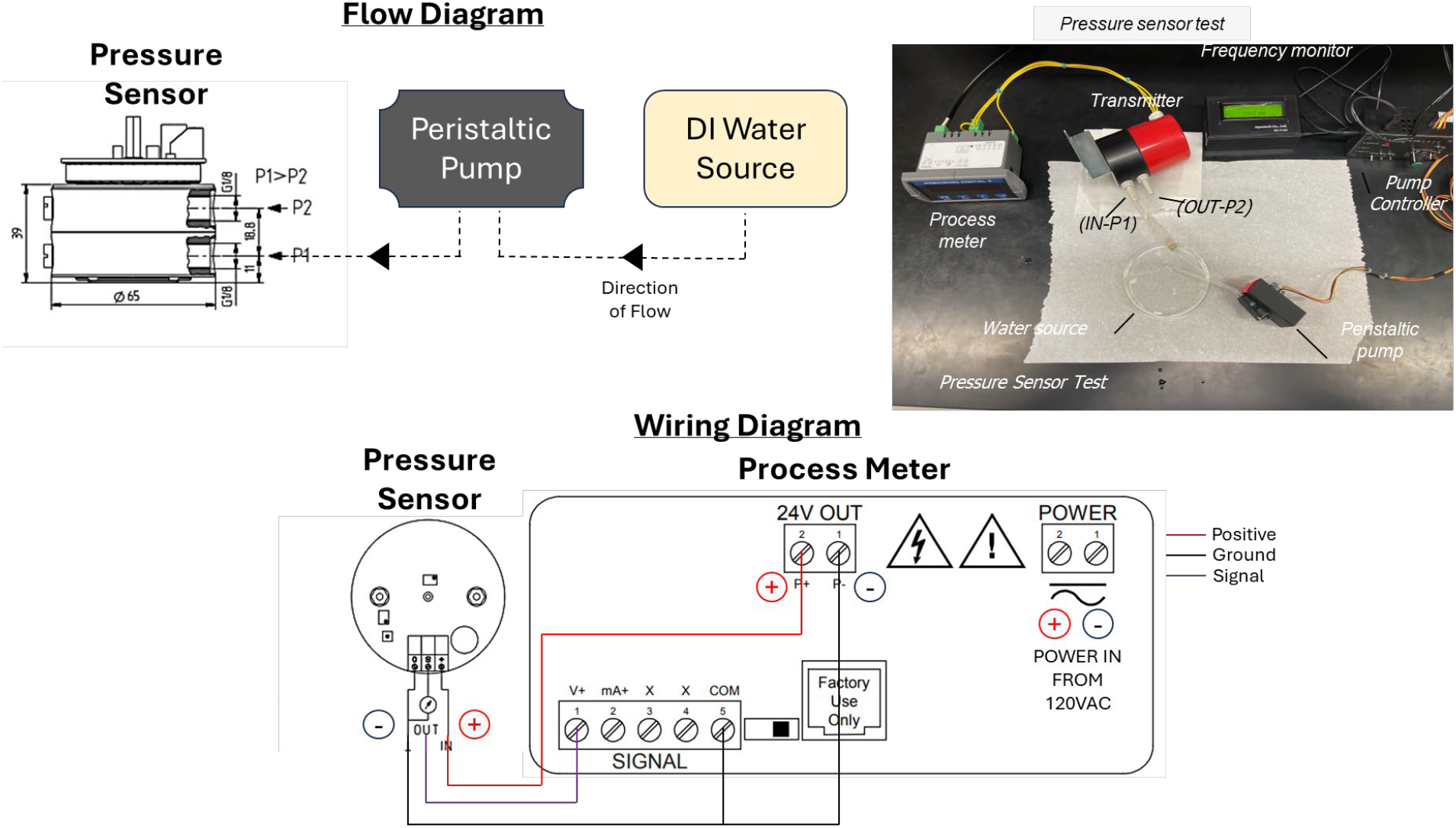
Schematic of connections, instrument parts, and materials to measure the pressure administered by the peristaltic pumps

**Movie 1**. Caption: Representative video file showing deformation of collagen gel with encapsulated fluorescent beads in chamber 2 when subjected to PUFFS at 0.33 Hz. (Captured at 7 frames per second)

**Movie 2**. Caption: Representative video file showing calcium signal propagation (right to left) across 3D MLO-Y4 networks in chamber 2 during PUFFS application at 0.33 Hz.

